# Retrograde transport of neurotrophin receptor TrkB-FL induced by excitotoxicity regulates Golgi stability and is a target for stroke neuroprotection

**DOI:** 10.1101/2024.10.29.620835

**Authors:** Gema María Esteban-Ortega, Margarita Díaz-Guerra

## Abstract

Excitotoxicity, aberrant function of survival pathways dependent on brain-derived neurotrophic factor (BDNF) and disruption of the Golgi complex are shared pathological hallmarks of relevant chronic and acute neurological diseases, including stroke. However, precise interdependence among these mechanisms is not completely defined, a knowledge essential to develop neuroprotective strategies. For ischemic stroke, a leading cause of death, disability and dementia, promising results have been obtained by interfering excitotoxicity, major mechanism of neuronal death in the penumbra area surrounding the infarct. We are exploring neuroprotection by promotion of survival cascades dependent on BDNF binding to full-length tropomyosin-related kinase B (TrkB-FL) receptor, which become aberrant after excitotoxicity induction. We have previously developed a blood–brain barrier (BBB) permeable neuroprotective peptide (MTFL_457_) containing a TrkB-FL sequence which efficiently prevents receptor processing induced by excitotoxicity and preserves BDNF-dependent pathways in a model of ischemia, where it efficiently decreases infarct size and improves neurological outcome after stroke. In this work, using cellular and animal models, we demonstrate that excitotoxicity-induced TrkB-FL downregulation is secondary to receptor endocytosis, receptor interaction with endosomal protein hepatocyte growth factor-regulated tyrosine kinase substrate (Hrs), retrograde transport to the Golgi and disruption of this organelle. Interestingly, peptide MTFL_457_ efficiently interferes TrkB-FL/Hrs interaction and receptor trafficking, processes required for excitotoxic Golgi fragmentation and TrkB-FL cleavage, demonstrating a central role for TrkB-FL in the control of Golgi stability. These results also suggest the potential of peptide MTFL_457_ to preserve function of this organelle and of critical neuronal survival pathways in stroke and, probably, other neurodegenerative diseases associated to excitotoxicity.

## Introduction

Stroke is a highly relevant disease affecting world population. It is currently a leading cause of death, disability and dementia, its burden expected to increase due to world population aging and higher prevalence of vascular risk factors. However, no effective therapies have yet reached the clinic. For ischemic stroke treatments are largely limited to mechanical or pharmacological blood flow restoration and only reach few patients. Therefore, we need to develop new safe and efficient treatments to reduce stroke impact. Disruption of cerebral blood flow by vessel occlusion generates an irreversibly damaged tissue, the infarct core, surrounded by a potentially recoverable “penumbra” area. However, this region may undergo secondary neuronal death, mainly due to overstimulation of the N-methyl-D-aspartate type of glutamate receptors (NMDARs) and consequent excitotoxicity [1], causing infarct expansion. Thus, current neuroprotective strategies are mostly directed to penumbra rescue through interference of excitotoxicity. Interestingly, this process is also associated to many other CNS disorders such as hypoglycaemia, epilepsy, acute trauma and neurodegenerative diseases (NDDs) [2], reinforcing the importance of excitotoxicity- targeted therapies to reduce neurological damage. NMDAR antagonists have failed as neuroprotectants due to their lack of specificity but promising results have been obtained targeting NMDAR-downstream signalling [3]. Thus, clinical-stage results have been obtained for acute ischemic stroke [4, 5] uncoupling NMDAR overactivation and neurotoxic signalling through dissociation of the ternary complex formed by GluN2B NMDAR-subunits, postsynaptic density protein-95 (PSD-95) and neuronal nitric oxide synthase (nNOS) [6]. This approach has leant on cell-penetrating peptides (CPPs), promising molecules for CNS treatment acting as carriers to drive therapeutic molecules across the blood–brain barrier (BBB) and plasma membrane [7]. We have also employed CPPs to investigate complementary approaches aimed to prevent aberrant excitotoxic downregulation of survival pathways, such as those regulated by full-length tropomyosin-related kinase B receptor (TrkB-FL) [8] or PSD-95 [9].

Promotion of neuronal survival is facilitated by brain-derived neurotrophic factor (BDNF) binding to its high affinity receptor TrkB-FL, increased tyrosine kinase (TK) activity and transphosphorylation of residues Y515 and Y816, anchor sites for adaptor proteins [10]. The result is the activation of interconnected cascades, including that regulated by phospholipase C © (PLC©) [10, 11] that, among other functions, activates prosurvival transcription factors (TFs) cAMP response-element binding protein (CREB) [12] and myocyte enhancer factor 2 (MEF2) [13, 14]. Their targets include genes encoding for TrkB [15, 16] and BDNF [17–19]. Upon neurotrophin binding, TrkB-FL and TrkB-T1, a truncated TK- deficient isoform considered a TrkB-FL dominant-negative neuronal receptor [20], are quickly and efficiently internalized [21, 22] forming signalling endosomes [22]. Recycling back to cell-surface is less efficient for TrkB-FL, relies on its TK activity, and is regulated by binding of the interdomain region separating transmembrane (TM) and TK sequences to endosomal protein hepatocyte growth factor-regulated tyrosine kinase substrate (Hrs) [22].

TrkB signalling becomes profoundly aberrant in stroke and NDDs [23, 24], pointing to receptor isoforms as therapeutic targets for neuroprotection in excitotoxicity-associated diseases. Three mechanisms contribute to TrkB dysregulation: 1) transcriptional regulation favouring TrkB-T1 expression over TrkB-FL [25, 26]; 2) TrkB-FL cleavage by calpain producing a truncated receptor alike to TrkB-T1 [26] and a cytosolic 32-kDa fragment (f32) [25] presenting TK activity and nuclear accumulation [27]; and 3) regulated intramembrane proteolysis (RIP) of both isoforms by concerted metalloproteinase (MP)/γ- secretase action releasing identical ectodomains which sequester BDNF [26]. In stroke, TrkB-FL regulation is mostly due to calpain-processing while RIP is only secondary. We have demonstrated before that TrkB-FL is depleted from neuronal surface early in excitotoxicity, and calpain and RIP processing require previous endocytosis [8], suggesting cleavage in intracellular compartments. To preserve BDNF/TrkB-FL survival cascades in excitotoxicity, we developed a HIV Tat-derived CPP (MTFL_457_) containing a TM/TK interdomain sequence able to maintain TrkB-FL in the cell-surface, away from the excitotoxicity-activated proteolytic machinery, secondarily preventing processing by calpain and MPs [8]. As a result, neuronal viability was preserved via a PLC©-dependent cascade initiating a feedback mechanism mediated by maintenance of downstream CREB and MEF2 promoter activities and increased expression of critical prosurvival proteins. This neuroprotective CPP might be relevant to human stroke therapy since it also prevents TrkB-FL downregulation in mouse ischemia where it efficiently reduces infarct size and neurological damage [8].

In this work, we demonstrate that TrkB-FL endocytosis induced by excitotoxicity is followed by interaction with Hrs and retrograde transport to Golgi apparatus (GA), in correspondence to organelle dispersion. This is important because GA disruption is considered a relevant hallmark of neuronal damage which precedes neurodegeneration [28]. Interestingly, MTFL_457_ inhibits TrkB-FL/Hrs interaction and blocks TrkB-FL retrograde transport, resulting in neuronal protection from organelle fragmentation and excitotoxicity. Thus, our results unveil a central role for TrkB-FL in GA stability and suggest that neuroprotective peptide MTFL_457_ could be relevant not only for treatment of stroke but also NDDs associated to excitotoxicity and GA disruption.

## Results

### MTFL_457_ preserves a pY816-TrkB-FL receptor at the cell-surface away from the proteolytic machinery secondarily activated by excitotoxicity

Previous results suggested that processing was not the primary mechanism of TrkB-FL regulation in excitotoxicity [8]. Analysis of receptor downregulation at early times of *in vitro* excitotoxicity showed maintenance of characteristic TrkB-FL labelling (Fig. 1Aa) up to 30 min of damage (Fig. 1A, b and c). NMDA treatment progressively decreased TrkB-FL signal (Fig. 1A, d and e), particularly in dendrites which presented focal swellings and varicosities (arrowheads) characteristic of excitotoxicity, preceding death [29]. For the longest treatment, there was also partial nuclear accumulation (Fig. 1Ae, inset), probably due to f32 translocation [27]. Changes in TrkB-FL levels were not statistically significant until 90 min (Fig. 1C) and were paralleled by f32 increase and concomitant to calpain activation, established by cleavage of its well-characterized substrate spectrin into breakdown products (BDPs, Fig. 1B). Downregulation of additional calpain substrates such as PSD-95 [9] and brain spectrin [30] was faster (Fig. 1B, D) and specific, other proteins such as neuronal specific enolase (NSE) not being significantly affected. The idea that TrkB-FL processing was secondary to other events was also supported by inhibition with dynasore of dynamin-dependent endocytosis which resulted in TrkB-FL stabilization in excitotoxicity (Fig. 1E, F), as previously demonstrated in ischemia [8]. Moreover, in contrast to control peptide MTMyc (Fig. 1G) which had no effects on TrkB-FL downregulation or neuronal viability [8], MTFL_457_ prevented progressive f32 accumulation and shedding of the highly glycosylated receptor ectodomain induced by NMDA (TrkB-ECD, Fig. 1H, I). Interestingly, this TrkB-derived peptide lacks the sequences presumably recognized by calpain and MPs and, therefore, must affect receptor processing in an indirect way. In fact, MTFL_457_ prevents pY816-TrkB-FL endocytosis and preserves this partially active receptor form at the cell-surface (Fig. 1J, K). In contrast, a brief NMDA treatment strongly induces receptor endocytosis in MTMyc-treated cultures, probably due to general triggering of endocytic processes by excitotoxicity [31, 32].

**Fig. 1.**
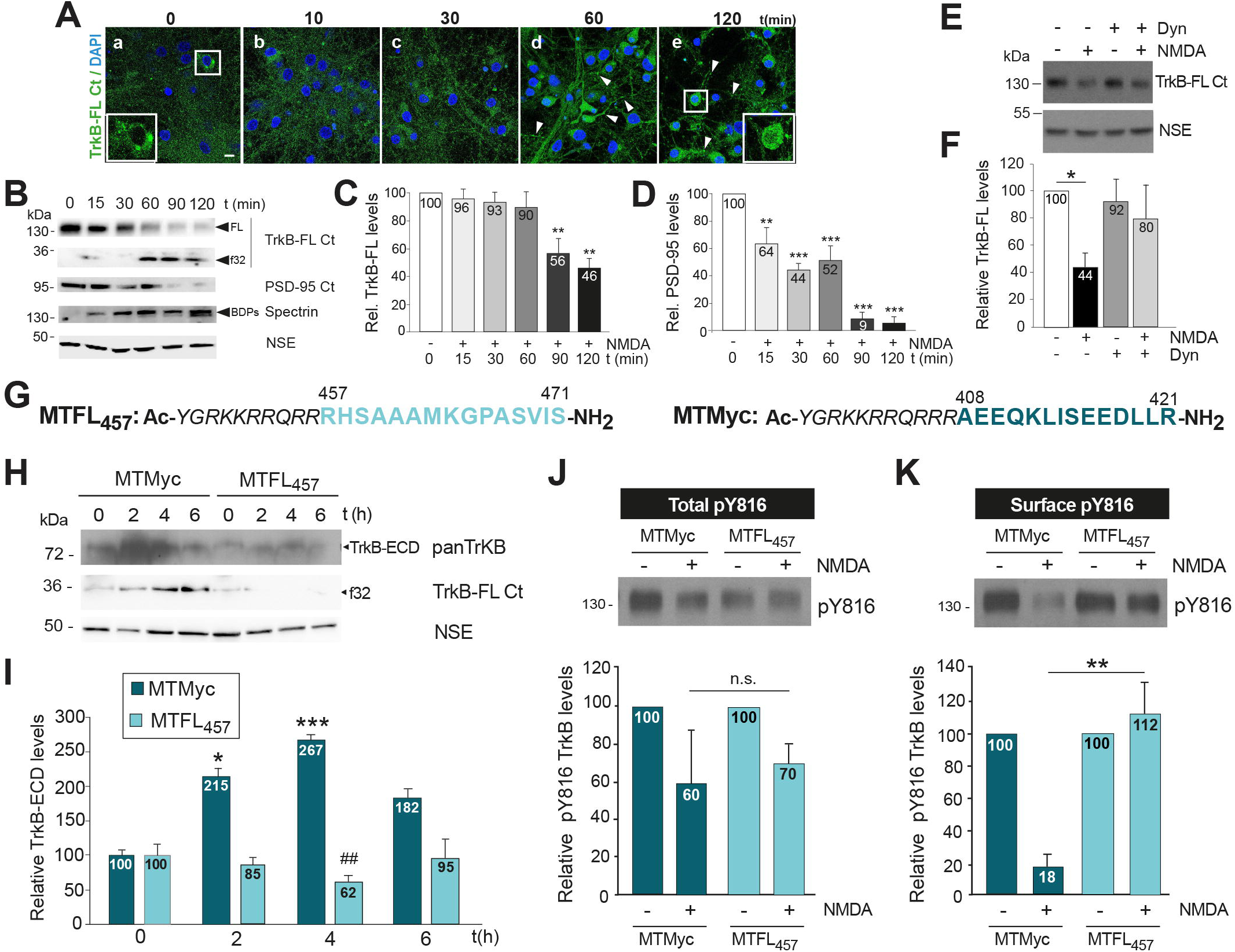
MTFL_457_ preserves a pY816-TrkB-FL receptor at cell-surface away from proteolytic machinery secondarily activated by excitotoxicity. **A-D** Kinetics of TrkB-FL downregulation. Primary cortical cultures were treated with NMDA (100 μM) and its coagonist glycine (10 μM), denoted as “NMDA”. Immunofluorescence (A) and immunoblot (B) were performed with a C-ter isoform-specific antibody (TrkB-FL Ct) recognizing full-length protein (FL) and intracellular fragment (f32). A TrkB-FL (green) and nuclei (blue), detected with DAPI. Arrowheads indicate varicosities in neuronal projections. Scale bar: 10 μm. Cell body details are shown for untreated cells or those with NMDA (120 min). B Decrease of TrkB-FL and f32 formation compared to PSD-95 downregulation, detected by a C-ter antibody (PSD-95 Ct). Calpain activation was established by accumulation of characteristic spectrin breakdown products (BDPs; 150 and 145 kDa). Neuron-specific enolase (NSE) was used as a loading control and for protein normalization. C, D Quantification of normalized TrkB-FL and PSD-95 levels relative to values in the absence of NMDA. Means ± SEM are represented. Statistical analysis was performed by one-way analysis of variance (ANOVA) test followed by a Bonferroni *post hoc* test (\**\*P* < 0.01, *** *P* < 0.001; *n =* 5). E, F Cultures were preincubated or not with dynasore (80 μM, 30 min) and treated with NMDA (2 h). Means ± SEM (*n =* 4), statistical analysis as above (*\*P* < 0.05). G Sequences of cell-penetrating neuroprotective (MTFL_457_) and control (MTMyc) peptides, which contain Tat aa 47–57 (italic) followed by, respectively, rat TrkB-FL (aa 457-471, light blue) or c-Myc (aa 408-421, dark blue) sequences. **H**, I Cultures were preincubated with peptides (25 μM, 30 min) followed by NMDA treatment. Culture media was analysed with antibodies directed to TrkB-FL extracellular domain (panTrkB), recognizing all isoform ECDs (TrkB-ECD), to investigate receptor shedding by MP activation. Analysis of total lysates with TrkB-FL Ct investigated TrkB-FL calpain-processing through f32 production. Means ± SEM (*n =* 7) of TrkB-ECD values are represented, and statistically analysed by ANOVA followed by a Bonferroni’s honestly significant difference test (*\*P* < 0.05, \*\*\**P* < 0.001, MTMyc + NMDA vs MTMyc; ^#*#*^*P* < 0.01, MTMyc vs MTFL_457_ at each time). **J, K** NMDA effect on total and cell-surface pY816-TrkB-FL levels. Cultures were preincubated with peptides as above, before a brief NMDA treatment (1 h) to avoid extensive receptor degradation, followed by biotin-labelling and sedimentation of membrane proteins, and comparison to corresponding total lysates. Means ± SEM (*n =*4) are represented and analysed by unpaired Student’s t-test (n.s. = not significant; \**\* P* < 0.01).

### MTFL_457_ neuroprotective effect is independent of TrkB-FL KFG domain

Regulation of TrkB-FL endocytosis in a pathological context such as excitotoxicity has not been characterized before. BDNF-induced endocytosis is controlled by TrkB-FL ubiquitination [33]. While controversial for TrkB [34], K460 ubiquitination inside a highly conserved KFG domain is important in traffic regulation of Trk receptors. MTFL_457_ contains K460 and surrounding residues and, therefore, it might potentially prevent TrkB ubiquitination and endocytosis induced by NMDA. In contrast, a peptide lacking the KFG sequence (MTFL_457_AAA, Fig. 2A) would not affect neuronal viability *in vitro* or *in vivo*. However, MTFL_457_ and MTF_457_AAA similarly prevented the significant decrease induced by NMDA in neuronal viability in MTMyc-preincubated cultures (Fig. 2B). For *in vivo* experiments, we used a model of microvascular photothrombosis (Fig. 2C) [35]. Permanent vessel occlusion and focal cortical infarcts were produced in motor and somatosensory areas of the ipsilateral hemisphere, visualized as TTC unstained regions (Fig. 2D). Infarct volumes in animals treated with MTF_457_ and MTF_457_AAA were very similar and significantly lower than values obtained with MTMyc, representing a 28% reduction (Fig. 2E). Therefore, the KFG sequence is not a requirement for MTFL_457_ ability to interfere *in vitro* and *in vivo* excitotoxicity.

**Fig. 2.**
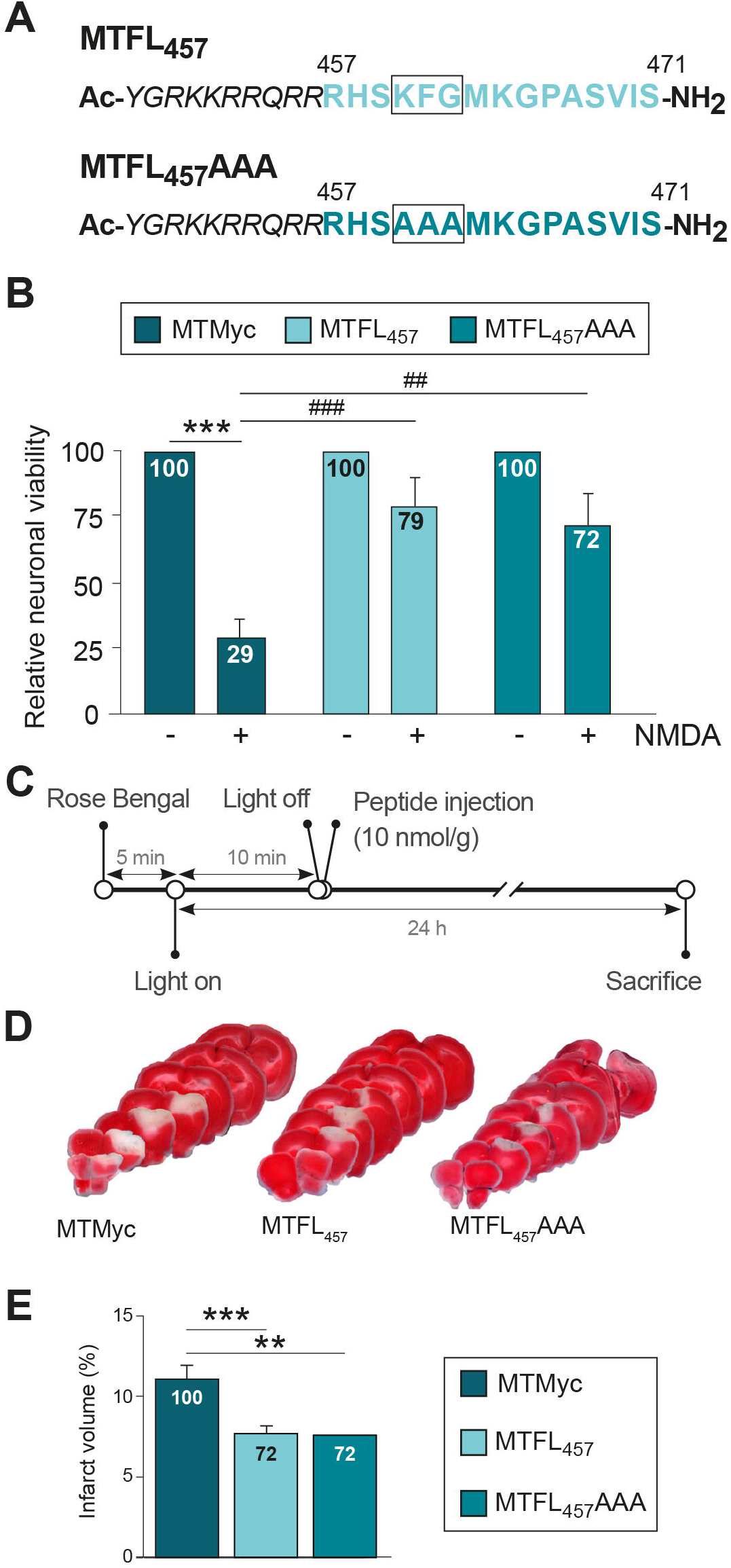
MTFL_457_ neuroprotective effect is independent of the TrkB-FL KFG domain. **A** Comparison of MTFL_457_ and MTFL_457_AAA sequences. The latter contains Tat aa 47–57 (italic) followed by the indicated rat TrkB-FL sequence but substituting the highly conserved KFG domain by AAA. **B** Effect of MTFL_457_AAA on neuronal viability in cultures preincubated with MTMyc, MTFL_457_ or MTFL_457_AAA as before, and treated with NMDA (2 h). Results obtained in the excitotoxic conditions were expressed relative to those obtained in the absence of NMDA. Means ± SEM (*n =* 5) are represented, and statistical analysis was performed by ANOVA followed by a Bonferroni test (\*\**\*P* < 0.001 for NMDA effect on MTMyc- treated cells; ##*#P* < 0.001 and #*#P* < 0.01 for CPP effect on NMDA-treated cells). **C** Timeline to analyse *in vivo* effects of MTMyc, MTFL_457_ or MTFL_457_AAA in the model of ischemia. Microvascular photothrombotic permanent damage was initiated in mice by cold-light irradiation (10 min) of a stereotaxically selected brain area after i.v. injection of photosensitive dye Rose Bengal as detailed in Material and Methods. CPPs (10 nmol/g) were retro-orbitally administered 10 min after damage initiation and animals were sacrificed 24 h later. **D** Representative 1 mm brain coronal slices stained with TTC corresponding to animals injected with MTMyc, MTFL_457_ or MTFL_457_AAA. **E** Infarct volume of CPP- injected animals expressed as a percentage of the hemisphere volume. Means ± SEM are given (*n =* 9). Differences were analysed by Student’s t-test (\**\*P* < 0.01, \*\**\*P* < 0.001). Infarct volume for MTFL_457_ and MTFL_457_AAA experimental groups are also expressed as a percentage of values obtained in animals injected with MTMyc.

### Effect of excitotoxicity in TrkB-FL interaction with endosomal protein Hrs and MTFL_457_ regulation

Abundance of membrane proteins at the cell-surface depends on integration of secretory and endolisosomal pathways which establishes the steady state proteome, susceptible to active remodelling depending on changing physiologic needs. Disruption of this balance is associated to pathological conditions, particularly diseases related to aging and neurodegeneration [36]. In BDNF response, TrkB-FL juxtamembrane intracellular region L453-N536 is involved in an interaction with Hrs, an early and late endosomal protein [37] regulating balance between receptor partial lysosomal degradation or recycling back to membrane [22, 37, 38]. We hypothesized that excitotoxicity might profoundly alter this balance normally regulated by Hrs.

To test this hypothesis, we first investigated a possible effect of excitotoxicity on Hrs levels and found no changes after brief (Fig. S1A) or prolonged (Fig. S1B, C) induction. Next, we analysed a possible TrkB-FL/Hrs interaction in basal or excitotoxic conditions. Colocalization was low in basal conditions (Fig. 3A) in correspondence to a Pearson correlation coefficient (PCC) below a threshold value of 0.50 (Fig. 3B). Excitotoxicity induced a progressive increase in colocalization (Fig. 3A), the PCC reaching a value of 0.56 after 60 min (Fig. 3B), when 83 ± 9% of neurons had a PCC ≥ 0.50 (Fig. 3C). We confirmed these results by coimmunoprecipitation with Hrs antibodies (Fig. 3D, E) or a generic TrkB antibody (panTrkB; Fig. S2). In cultures briefly treated with NMDA or BDNF, used as an internal control, we confirmed sustained Hrs levels in total extracts and comparable immunoprecipitation by Hrs antibody (Fig. 3D). In contrast, although a significant TrkB-FL decrease was induced by NMDA, similar levels were coimmunoprecipitated by the Hrs antibody (Fig. 3E). The difference between relative TrkB-FL levels in total extracts or Hrs-coimmunoprecipitated was statistically significant in excitotoxicity. As described, treatment with BDNF increased TrkB-FL/Hrs interaction [22]. For experiments using panTrkB, we confirmed TrkB-FL precipitation with isoform-specific antibody TrkB-FL Ct (Fig. S2A), finding a reduction induced by excitotoxicity in receptor levels in correspondence to that of total TrkB-FL. However, no parallel decrease was observed in coimmunoprecipitated Hrs (Fig. S2B).

**Fig. 3.**
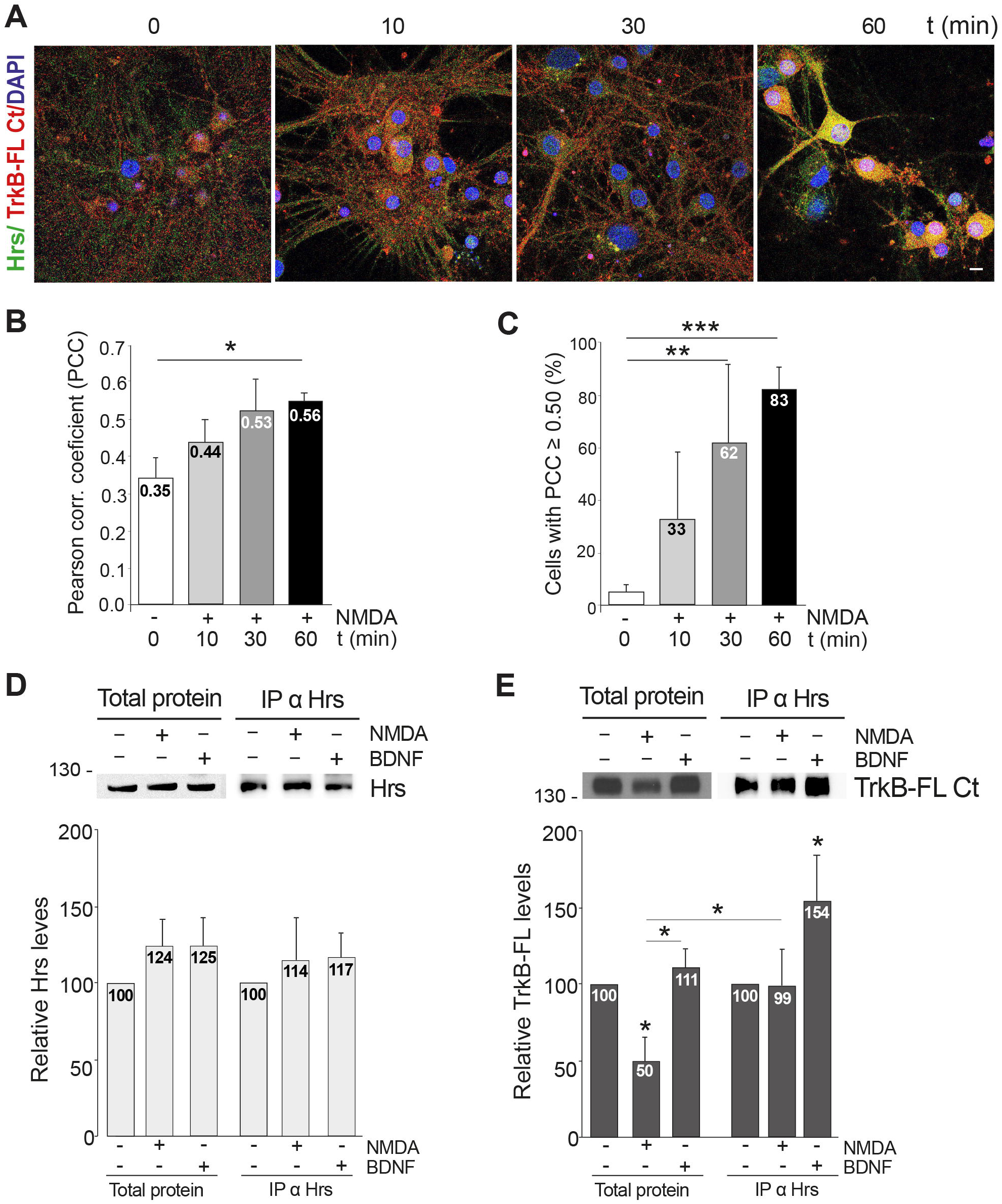
Effect of excitotoxicity in TrkB-FL interaction with endosomal protein Hrs. A-C. Analysis by immunofluorescence of TrkB-FL/Hrs colocalization. **A** Cortical neurons were treated with NMDA for the indicated times and analysed with antibodies specific for Hrs (green) and TrKB-FL Ct (red), nuclear staining being performed with DAPI (blue). Representative images obtained by confocal microscopy correspond to single sections and show the fused channels. Scale bar: 10 μm. **B** Mean values ± SEM of Pearson correlation coefficient (PCC; *n =* 3). For each independent experiment, a minimum of 80 different neurons were analysed. Statistical analysis was performed using a generalized linear model followed by a *post-hoc* LSD test (*\*P* < 0.05). **C** Percentage of cells showing at different times of NMDA treatment a PCC ≥ 0.50, generally considered as a threshold for protein colocalization. Statistical analysis was performed as before (\**\*P* < 0.01; \*\**\*P* < 0.001; *n =* 3). **D, E** Analysis by immunoprecipitation of TrkB-FL/Hrs interaction. Neuronal cultures were treated with NMDA (100 μM) or BDNF (100 ng/ml) for 30 min and compared to untreated cultures. Immunoprecipitation was performed with the Hrs antibody and the immunoprecipitated proteins (IP) were analysed by immunoblot, using the same antibody **(D)** or TrkB-FL Ct **(E)**. Total protein lysates were analysed in parallel to the immunoprecipitated proteins. Mean values ± SEM (*n =* 5) of Hrs and TrkB-FL levels in NMDA or BDNF-treated cultures relative to the untreated cells is represented both for total lysates or Hrs-immunoprecipitated proteins. Statistical analysis was performed using a generalized linear model followed by a *post-hoc* LSD test (*\*P* < 0.05).

MTFL_457_ contains TrkB-FL residues inner to the region involved in Hrs interaction. Therefore, we characterized a possible peptide effect on TrkB-FL/Hrs colocalization (Fig. 4). In MTMyc-cultures, excitotoxicity induced an increase in colocalization similar to that described in peptide absence (Fig. 4A, B). In contrast, colocalization was not affected by excitotoxicity in the presence of MTFL_457_. Coimmunoprecipitation experiments confirmed immunoprecipitation of equivalent Hrs amounts (Fig. 4C) while TrkB-FL/Hrs interaction was favoured by excitotoxicity in MTMyc-cultures (Fig. 4D). In contrast, although TrkB-FL was stabilized by MTFL_457_ action, excitotoxicity induced a significant decrease in interaction. We conclude that, from early times of NMDAR overactivation, Hrs interaction is promoted and regulates TrkB-FL fate. Disruption by MTFL_457_ of TrkB-FL/Hrs interaction might secondarily prevent excitotoxicity-induced TrkB-FL proteolysis, resulting in neuroprotection.

**Fig. 4.**
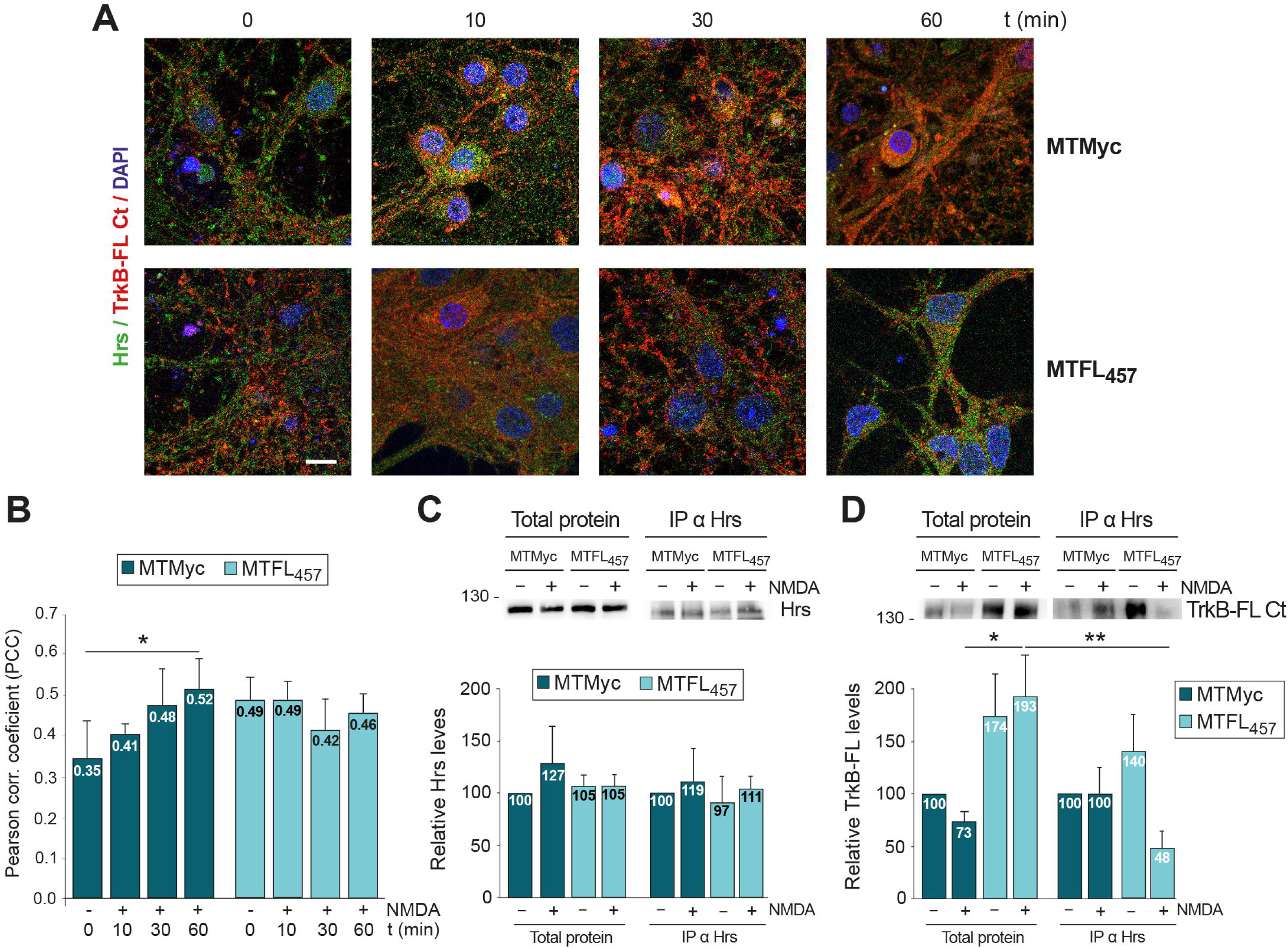
Regulation of TrkB-FL/Hrs interaction induced by excitotoxicity by MTFL_457._ **A** Cortical neurons were preincubated with MTMyc and MTFL_457_ (25 μM, 30 min) and treated with NMDA for the indicated times. Cells were analysed with antibodies for Hrs (green) and TrKB-FL Ct (red), together with DAPI staining (blue). Representative images obtained by confocal microscopy correspond to single sections and show the fused channels. Scale bar: 10 μm. **B** Mean values ± SEM of PCC (*n =* 3). For each independent experiment, a minimum of 80 different neurons were analysed. Statistical analysis was performed using a generalized linear model followed by a *post-hoc* LSD test (*\*P* < 0.05). **C, D** Analysis by immunoprecipitation of TrkB-FL/Hrs interaction. Cultures preincubated with CPPs as above were treated with NMDA for 30 min and compared to untreated cultures. Immunoprecipitation was performed with the Hrs antibody and the immunoprecipitated proteins (IP) were analysed by immunoblot, using the same antibody **(C)** or TrkB-FL Ct **(D)**. Total protein lysates and immunoprecipitated proteins were analysed in parallel. Mean values ± SEM (*n =* 3) of Hrs and TrkB-FL levels relative to those found in cells preincubated with MTMyc and without NMDA is represented. Statistical analysis was performed using a generalized linear model followed by a *post-hoc* LSD test (*\*P* < 0.05, \**\*P* < 0.01).

### Effect of excitotoxicity in TrkB-FL transport to the GA and MTFL_457_ regulation

Retrograde GA protein transport allows some cargoes to avoid lysosomal degradation, although they are still susceptible to proteolysis in that organelle. Thus, TrkB/NMDAR shared effector Kidins220 [39] undergoes endocytosis early after excitotoxicity, followed by GA transport and calpain-dependent degradation [40]. Accordingly, we decided to also characterize TrkB-FL GA trafficking after brief NMDA treatments by immunofluorescence with TrkB-FL Ct and a GM130 antibody recognizing this Golgi matrix protein, central to organelle preservation, protein glycosylation and vesicle transport. In basal conditions, GM130 showed a characteristic GA distribution in front of apical dendrites and a labelling pattern as flat cisternae (Fig. 5A) together with strong TrkB-FL presence (Fig. 5B) as described [41]. NMDA treatment further increased colocalization (Fig. 5B) causing TrkB-FL/GM130 colocalization in most neurons (Fig. 5C). Similar results were obtained in cultures preincubated with MTMyc (Fig. 5D-F) while TrkB-FL distribution was maintained in neurons preincubated with MTFL_457_. These results suggest that, early in excitotoxicity, a TrkB-FL fraction is recruited towards the GA before processing, and peptide MTFL_457_ can prevent this situation.

**Fig. 5.**
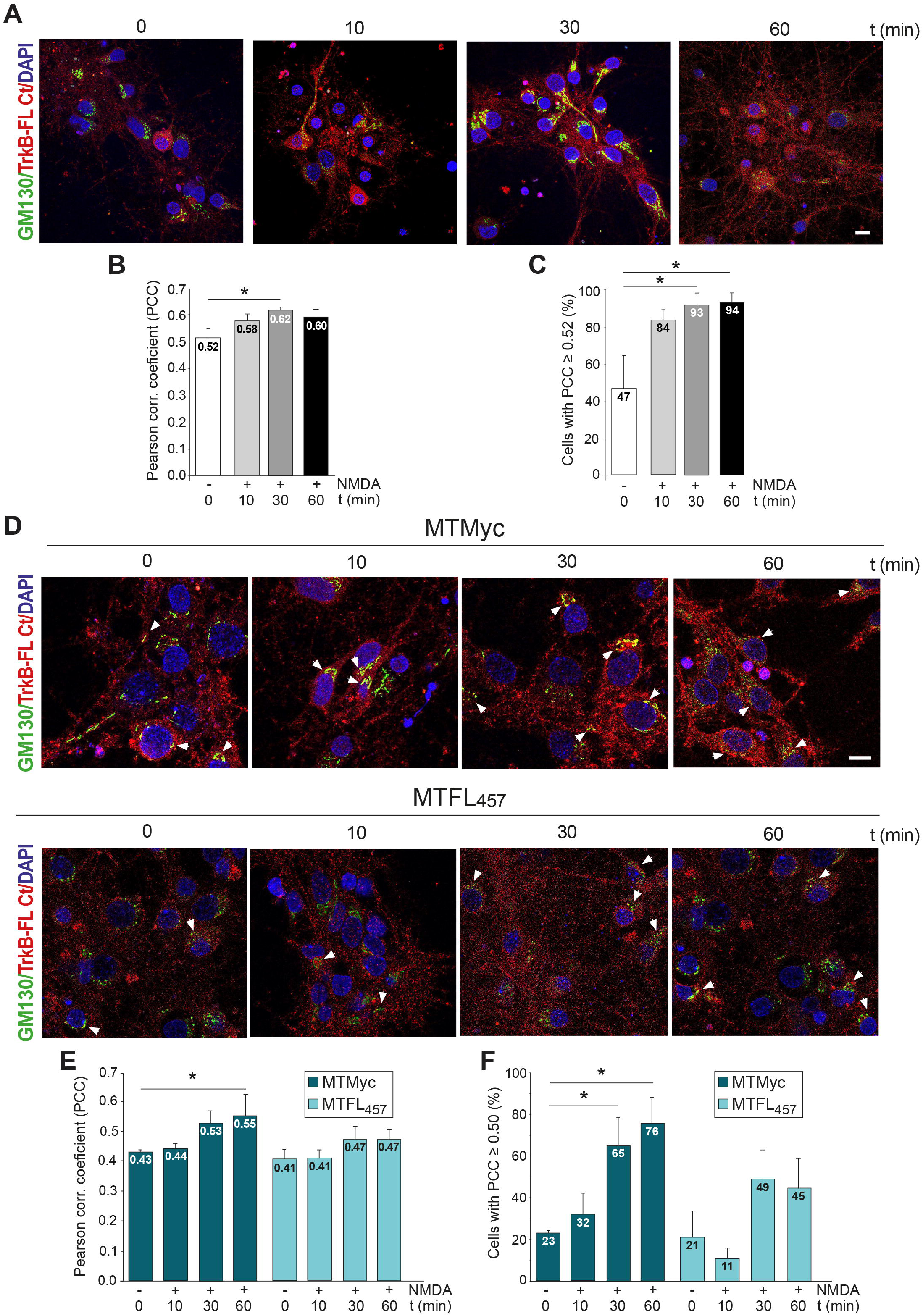
Effect of excitotoxicity in TrkB-FL transport to the Golgi complex and regulation by MTFL_457_ action. Cortical neurons were treated with NMDA for the indicated times and analysed by immunofluorescence with antibodies specific for the Golgi matrix protein GM130 (green) and TrkB-FL Ct (red), nuclear staining being performed with DAPI (blue). Excitotoxicity was induced in the absence of peptides **(A-C)** or after preincubation with MTMyc and MTFL_457_ (25 μM, 30 min) **(D-F)**. **A, D** Representative images obtained by confocal microscopy corresponding to single sections, showing independently the three channels or the fused image. Scale bar: 10 μm. **B, E** Mean PCC values ± SEM (*n =* 4) for TrkB-FL and GM130 colocalization. **C, F** Percentage of cells showing at the different times of treatment a PCC ≥ 0.52 **(C)**, the value obtained for TrkB-FL/GM130 colocalization in basal conditions in the absence of peptide, or ≥ 0.50 **(F)**. For each independent experiment, a minimum of 80 different neurons were analysed. The statistical analysis was carried out using the ANOVA test followed by Bonferroni test (*\*P* < 0.05).

### Effect of *in vitro* excitotoxicity on GA stability and MTFL_457_ regulation

Golgi fragmentation and dispersal precede neuronal death triggered by excitotoxicity and other types of insults [42]. Thus, we analysed GA stability at early times of *in vitro* excitotoxicity and how GA disruption might be affected by neuroprotective peptide MTFL_457_. Differently from condensed flattened cisternae found in untreated neurons, excitotoxicity induced a high degree of GA fragmentation driving the formation of scattered particles (Fig. 6A). In a representative experiment, we observed a gradual and parallel decrease with treatment of mean fluorescence intensity and area of detected structures (Fig. 6B). Differences in area (Fig. 6C), mean intensity (Fig. 6D) and integrated density (Fig. 6E) were statistically significant. Finally, particle circularity showed a tendency to increase as a consequence of fragmentation (Fig. 6F). These results confirmed GA disruption by excitotoxicity with kinetics similar to TrkB-FL retrograde transport. GA accumulation of TrkB-FL due to retrograde transport induced by excitotoxicity might promote organelle fragmentation and, therefore, prevention of receptor traffic by MTFL_457_ might also protect neurons from GA fragmentation. In fact, a remarkable decrease of GA fragmentation was observed after MTFL_457_ pre-treatment compared to MTMyc (Fig. 7A). In a representative experiment, MTFL_457_-treated neurons presented higher values of particle intensity and area relative to those preincubated with MTMyc (Fig. 7B). Accordingly, increased circularity was induced by NMDA in MTMyc-treated neurons, mainly associated to small particles (Fig. S3, left panel), a more dispersed distribution observed in MTFL_457_-treated cells (Fig. S3, right panel). Significant differences in particle area (Fig. 7C) and integrated density (Fig. 7D) found in MTMyc-cultures subjected to excitotoxicity were lost with MTFL_457_. Above results demonstrate that GA fragmentation is an early event induced by excitotoxicity preceding nuclear condensation or fragmentation which results in loss of a condensed GA structure in favour of more rounded and smaller particles, possibly vesicular fragments. MTFL_457_, a neuroprotective peptide preventing TrkB-FL retrograde transport and degradation in excitotoxicity, is also capable of partially maintaining GA integrity. Therefore, neurotrophin receptor TrkB-FL might play a role in the regulation of Golgi disruption induced by neurodegeneration.

**Fig. 6.**
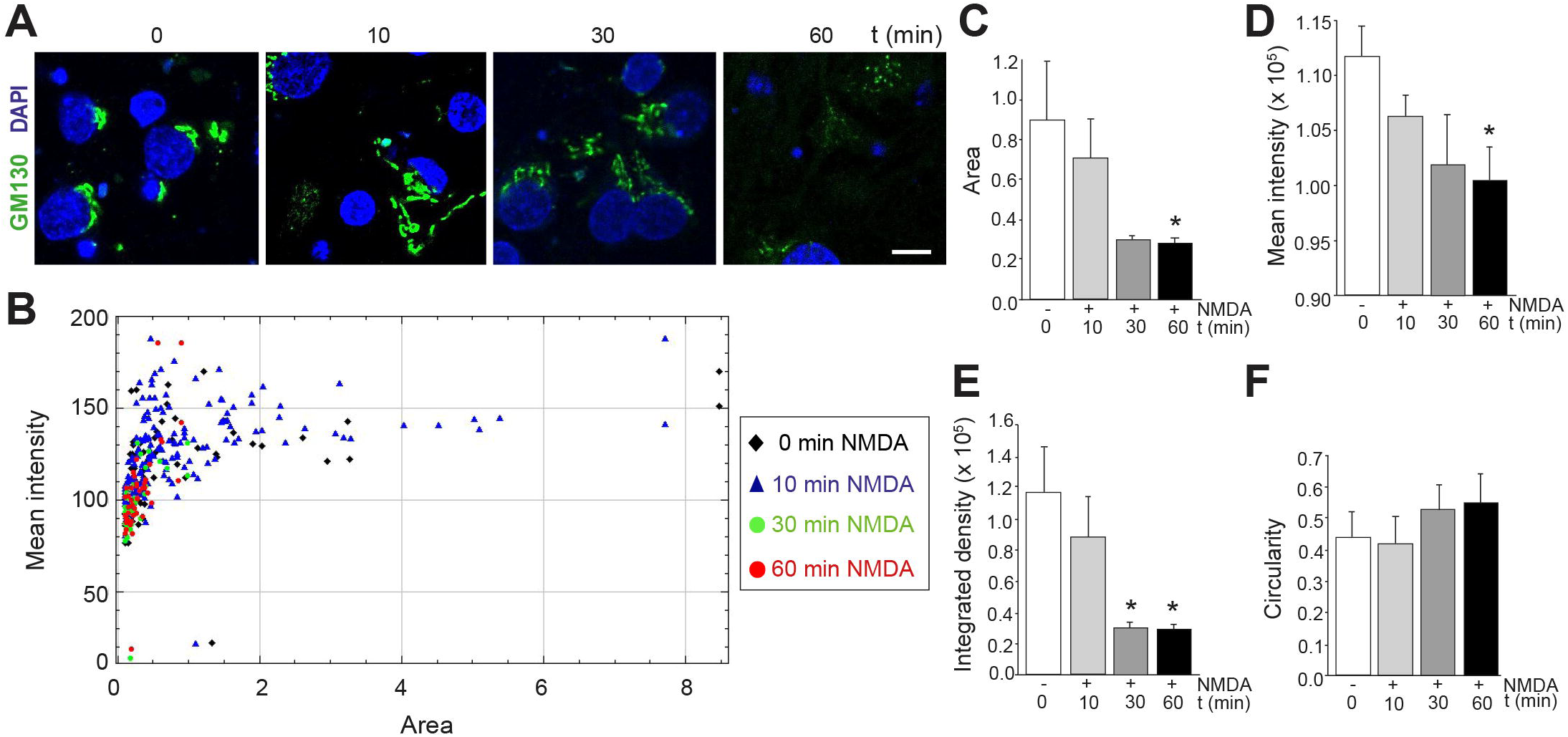
Effect of *in vitro* excitotoxicity on GA stability. Cortical neurons were treated with NMDA for the indicated times and analysed by immunofluorescence with GA specific antibody (GM130, green) together with DAPI nuclear staining (blue). **A** Representative images obtained by confocal microscopy corresponding to single sections. Scale bar: 10 μm. **B** Representation of the area versus the mean fluorescence intensity for each of the GA particles detected in representative images corresponding to the different times of NMDA treatment. Mean values ± SEM (*n =* 6) of the particles area **(C)**, mean fluorescence intensity **(D)**, integrated density **(E)** or circularity **(F)**, 0 value indicating absence of circularity, and 1 total circularity. An average of 80 particles were quantified per each independent experiment analysed. In all cases, the statistical analysis was carried out using ANOVA followed by the Bonferroni test (*\*P* < 0.05).

**Fig. 7.**
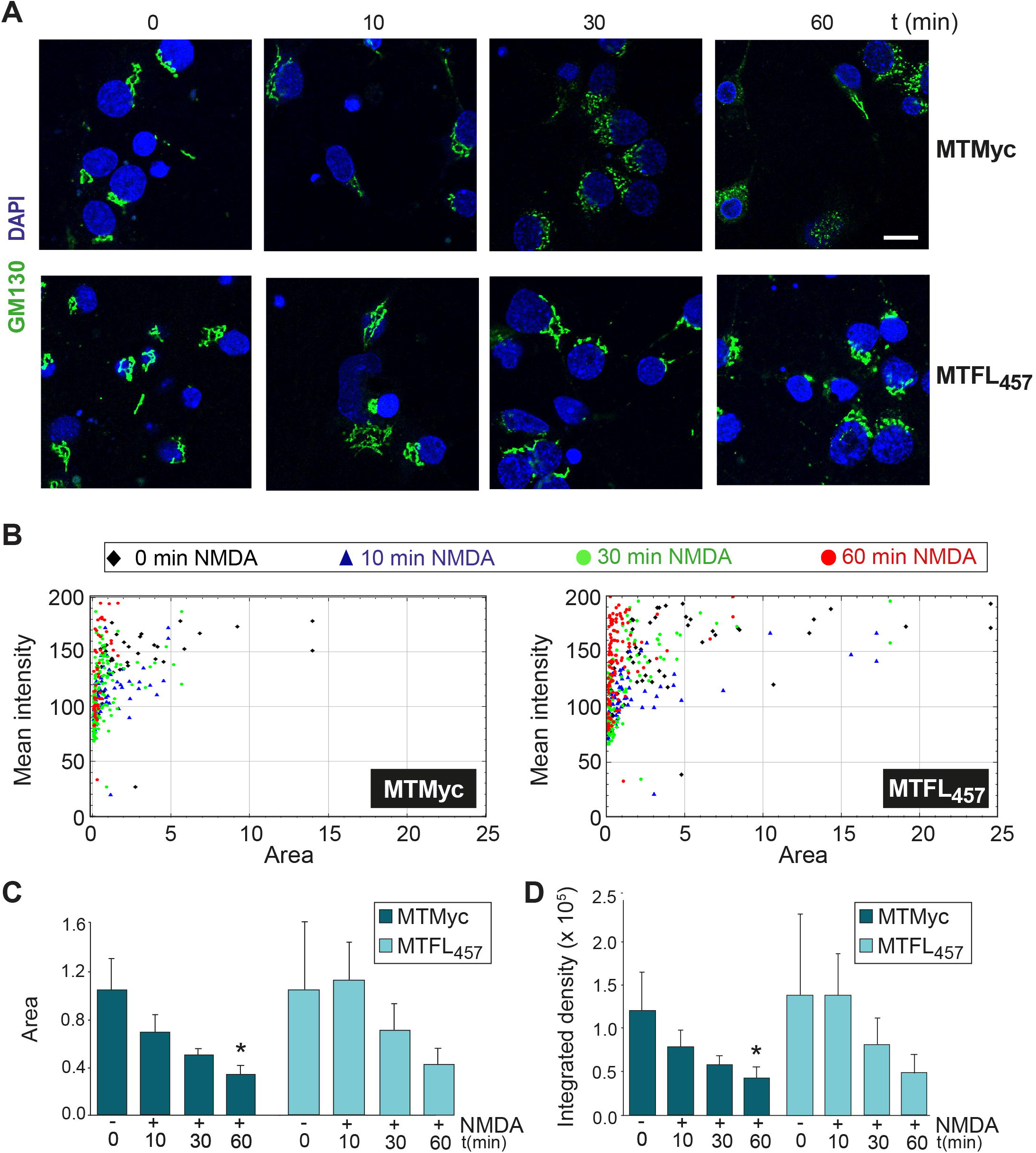
**Regulation of excitotoxicity-induced GA disruption by peptide MTFL_457._** Cortical neurons preincubated with MTMyc and MTFL_457_ (25 μM, 30 min) and treated with NMDA for the indicated times were analysed by immunofluorescence with GA specific antibody (GM130, green) and DAPI (blue). **A** Representative images obtained by confocal microscopy corresponding to single sections. Scale bar: 10 μm. **B** Representation of the area versus the mean fluorescence intensity for each of the GA particles detected in representative images corresponding to cultures preincubated with MTMyc (left panel) or MTFL_457_ (right panel), treated with NMDA for the indicated times. **C** Mean values ± SEM (*n =* 5) of the particles area. Statistical analysis was performed using ANOVA followed by the Bonferroni test (*\*P* < 0.05). **D** Mean values ± SEM (*n =* 5) of the integrated density. An average of 80 particles were quantified per each independent experiment analysed. Results were analysed using a generalized linear model followed by a *post-hoc* LSD test (*\*P* < 0.05).

### Effect of brain ischemia on GA stability and MTFL_457_ regulation

MTFL_457_ is efficiently delivered to the brain cortex after i.v. administration and, in the model of permanent ischemia [35, 43], has important effects on TrkB-FL downregulation, infarct size and neurological damage [8]. We do not know if GA disruption is also associated to neurodegeneration in this model of damage, characterized by a relatively narrow ischemic penumbra [44], or the possible effect of MTFL_457_. In a different model of transient ischemia, damage to this organelle was shown in penumbral neurons by mechanisms not well defined [45]. We analysed emerging infarcts produced 5 h after photothrombosis, well before infarct stabilization at 24 h of damage [26], comparing neurodegeneration in three cortical areas: the contralateral hemisphere, the infarct core or infarct peripheral areas. Degenerating neurons were identified using specific marker Fluoro-Jade C (FJC) or immunohistochemistry with SNTF antibody, labelling cells having overactivated calpain (Fig. 8A). We observed no signs of neurodegeneration in contralateral tissue in contrast to the ischemic core, characterized by high levels of FJC and SNTF staining, an intermediate situation represented by the peripheral ischemic tissue. GA analysis with mouse GM130 antibody was hampered by early BBB breakage after ischemia [46] and leakage of serum proteins, including immunoglobulins, to ischemic brain. This results in high backgrounds and heavy staining of blood vessels in the ischemic core by secondary anti-mouse antibodies (Fig. S4). Difficulty to use the same experimental conditions for GA visualization in ischemic and non-ischemic tissue, forced internal comparisons in contralateral (Fig. S5A) or ischemic peripheral areas (Fig. S5B). In non-ischemic conditions, we observed a GA distribution similar to that previously described in cultured neurons and no peptide effect (Fig. S5A). Interestingly, in the peripheral ischemic tissue of MTFL_457_-treated animals, we found a GA staining similar to that of the contralateral region together with lower backgrounds (Fig. S5B), probably due to reduced BBB damage. Finally, coronal sections were preincubated with an anti-mouse IgG (Fab specific) before GM130 immunodetection to improve GA visualization (Fig. 8B, C). We observed a fragmented and dispersed GA in the peripheral ischemic tissue of MTMyc-treated animals, in contrast to those injected with MTFL_457_ (Fig. 8C) which again showed an organelle distribution resembling that of the contralateral region (Fig. 8B). These results suggest that TrkB-FL retrograde transport and processing might be part of a mechanism regulating GA fragmentation associated to stroke and, consequently, strategies blocking receptor traffic and promoting GA stability could improve organelle function and contribute to neuronal survival after an ischemic insult.

**Fig. 8.**
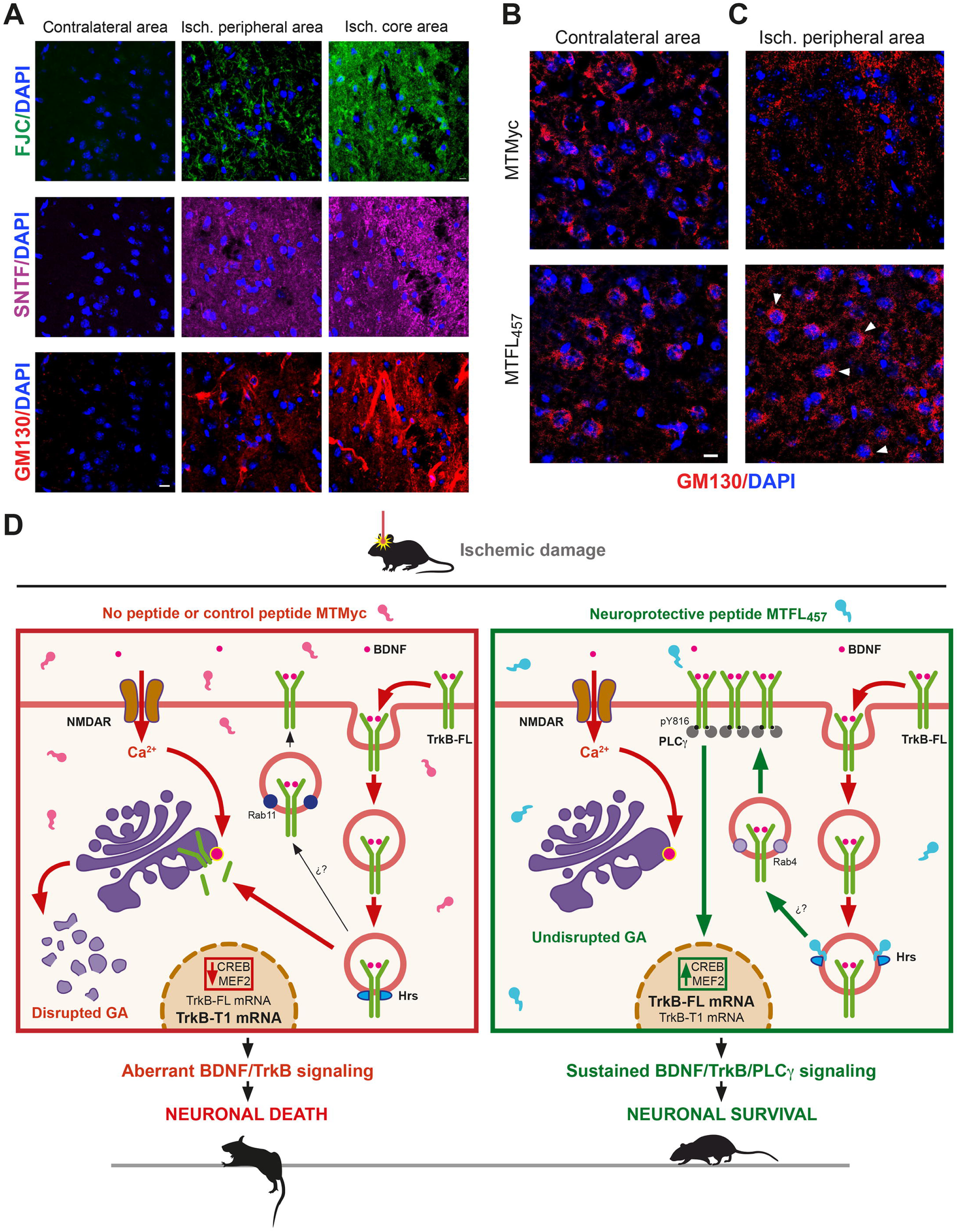
Effect of brain ischemia on GA stability and MTFL_457_ regulation and proposed model. **A** Strong association of neuronal degeneration and leakage of serum proteins after ischemic damage. Immunohistochemistry of brain coronal sections prepared from animals sacrificed 5 h after insult were performed with an antibody recognizing a calpain-generated neoepitope in spectrin N-terminal fragment (SNTF, magenta), which labels cells where this protease is overactive, and the mouse antibody recognizing GM130 (red). Neurodegeneration was also established by Fluoro-Jade C (FJC) staining (green). Three different areas of tissue were compared: the ischemic core, an area peripheral but close to the infarct core and the equivalent area of the contralateral hemisphere. Leakage of mouse immunoglobulins due to early BBB breakage after the ischemic insult, detected by the secondary anti-mouse antibody (see Fig. S5A), was observed in the neurodegenerating tissue and strongly interfered GM130 detection. **B, C** GM130 staining after preincubation of coronal sections with an anti-mouse IgG (Fab specific) antibody to improve detection. Animals were retro-orbitally injected with peptides MTMyc or MTFL_457_ (10 nmol/g) 10 min after damage initiation and sacrificed 5 h later. Comparison of the contralateral **(B)** or the ischemic peripheral areas **(C).** Representative images correspond to single sections. Scale bar: 10 µm. **D** Model of TrkB-FL regulation in excitotoxicity and MTFL_457_ action. Endocytosis of neurotrophin receptor TrkB-FL is promoted by excitotoxicity in neurons treated with control peptide MTMyc or without treatment (left panel). In endosomes, TrkB-FL interacts with protein Hrs and is retrogradely transported to the AG, where activation of organelle-associated proteinases would be responsible of receptor processing by calpain and RIP. Although there might be partial recycling back to the membrane by mechanisms similar to those found after BDNF activation, there is a strong decrease of BDNF/TrkB-FL signalling and CREB/MEF2 promoter activities, causing transcriptional changes that favour neuronal death. In parallel, the GA is disrupted, a hallmark common to many NDDs. Neuroprotective peptide MTFL_457_ interferes TrkB-FL/Hrs interaction induced by excitotoxicity, receptor retrograde transport and processing, as well as GA fragmentation (right panel). We propose that interference by MTFL_457_ of TrkB-FL/Hrs interaction might favour rapid recycling back to the membrane, similar to that of isoform TrkB-T1, sustained BDNF/TrkB-FL/PLC© signalling and promotion of neuronal survival.

## Discussion

We demonstrate here that neurotrophin receptor TrkB-FL plays a key role in GA fragmentation concomitant to stroke neurodegeneration, a discovery that encourages the development of innovative strategies for neuroprotection in ischemia and other NDDs associated to excitotoxicity [2] and GA disruption [28]. Excitotoxicity provokes a rapid decrease of cell-surface TrkB-FL and GA recruitment, processes preceding and necessary for organelle fragmentation (Fig. 8D, left panel). The GA is a highly dynamic organelle essential for functions including trafficking/sorting of membrane and secretory proteins, or glycosylation of lipids and proteins. Different physiological and pathological conditions modify GA structure. In mitotic cells, a reversible organelle disassembly is required for mitosis entry [47]. In contrast, in post-mitotic neurons, the GA is considered a sensor of stress signals in death cascades [42] and fragmentation precedes death induced, among others, by increased neuronal activity [42, 48]. Accordingly, GA disruption is an early event preceding further degeneration in stroke [45] and excitotoxicity-associated NDDs [2], including Alzheimer [49], Parkinson [50], Huntington [51] or amyotrophic lateral sclerosis [52]. Despite their critical consequences in neurons, the mechanisms involved in GA damage are poorly described. In line with our results, Golgi fragmentation is also induced by accumulation of autoactivated TrkB-FL in the ER–Golgi intermediate compartment due to heterologous overexpression or transactivation of endogenous receptors [53]. Consequently, increased TrkB-FL presence in the GA might have a regulatory role in damage-associated organelle fragmentation.

TrkB-FL intracellular traffic in a pathological situation such as excitotoxicity has not been previously characterized. Cell-surface receptor levels are strongly and rapidly decreased by excitotoxicity-induced endocytosis, similarly to GluN2B-containing NMDARs [54] or Kidins220 and GluN1-NMDAR subunits, retrogradely transported to the GA [40]. Likewise, internalized TrkB-FL is transported to this organelle in excitotoxicity (Fig. 8D, left panel), further increasing the significant receptor pool already present in basal conditions. Such an increase might constitute a stress signal promoting GA disruption as part of a cell death program. TrkB-FL retrograde transport requires Hrs interaction, suggesting that this endosomal protein regulates receptor trafficking in response to BDNF [38] and also NMDA. Differently from the physiological conditions, where Hrs balances TrkB-FL Rab11-dependent slow recycling back and lysosomal degradation, Hrs binding in excitotoxicity would favour receptor retrograde transport in detriment of lysosomal degradation. Secondary to receptor endocytosis and retrograde transport induced by excitotoxicity, TrkB-FL is processed by MPs/©-secretases and predominantly calpain [26], an effector of Ca^2+^-overload central to stroke and other acute or chronic pathologies likewise associated to excitotoxicity [30]. In addition to TrkB-FL [25], calpain cleaves different components of NMDAR-complexes critical to neuronal survival and function: scaffolding protein PSD-95 [9, 55], GluN2-NMDAR subunits [55] or Kidins220 [39, 56], which is similarly processed after GA transport [40]. Protease activation can occur close to plasma and endosomal membranes [57], high [Ca^2+^] microdomains or subcellular compartments such as GA membrane [58, 59], nucleus or mitochondria [60]. Excitotoxicity might trigger activation of GA-associated proteases and proteolysis of cargoes transported to this organelle, such as TrkB-FL (Fig. 8D, left panel). Altogether, these data unveil excitotoxicity-induced endocytosis, retrograde transport and processing of prosurvival proteins as relevant targets for neuroprotection [61].

Brain-accessible peptide MTFL_457_ has been critical to establish the importance of TrkB-FL retrograde transport for GA disruption and neurodegeneration in excitotoxicity. In addition to TrkB-FL endocytosis, MTFL_457_ prevents Hrs interaction, retrograde transport, organelle fragmentation and TrkB-FL proteolysis (Fig. 8D, right panel), underscoring the importance of residues 457-471 for control of receptor intracellular traffic induced by excitotoxicity. One possibility that deserves further investigation is promotion of TrkB-FL Rab4-dependent fast recycling, similar to that of TrkB-T1 [22], by interference of TrkB-FL/Hrs interaction. Anyway, a partially active pY816-TrkB-FL is preserved at the cell-surface, which supports neuronal viability via a PLC©-dependent mechanism probably mediated by maintenance of CREB and MEF2 activities (Fig. 8D, right panel). These TFs start a feedback mechanism favouring expression of critical prosurvival mRNAs and proteins in excitotoxicity [8]. These results resemble those obtained with classical and fast-acting antidepressants (ADs), which directly bind TrkB-FL dimers and mediate their effects on neuroplasticity by stabilizing a receptor conformation that facilitates membrane retention, synaptic location and BDNF accessibility [62]. Moreover, AD action depends on Y816-TrkB-FL phosphorylation and PLC© signalling, and results in increased CREB phosphorylation which mediates neuronal plasticity [63].

Neuroprotective peptide MTFL_457_ could be important for stroke treatment in humans because, in a preclinical stroke model, not only counteracts TrkB-FL downregulation, efficiently decreases infarct size, and improves neurological outcome [8] but also impairs GA fragmentation. In addition, MTFL_457_ might greatly enhance the efficiency of therapies for neurological and psychiatric diseases based on the use of BDNF or small size analogues [24] which are challenged by TrkB-FL downregulation and GA damage induced by excitotoxicity.

## Materials and methods

### Experimental models

Animal procedures were accomplished in agreement with European Union Directive 2010/63/EU and approved by ethics committees from CSIC and Comunidad de Madrid (Ref PROEX 276.6/20). The housing facilities were approved by Comunidad de Madrid (# ES 280790000188) and comply with official regulations. All efforts were made to minimize animal suffering and reduce number of sacrificed animals which had a standard health and immune status and were supervised daily by professional caretakers. Male mice were kept in groups (< 6) in standard individually ventilated cages, while 1-2 pregnant rats were maintained in standard cages, both types containing bedding and nesting material. Animals were under controlled lighting conditions (12 h light cycles), relative humidity and temperature, irradiated food and water provided *ad libitum*.

### Mouse model of ischemia by photothrombosis

Permanent focal ischemia was induced in the cerebral cortex of adult male Balb/cOlaHsd mice [25-30 g; 8-12 weeks of age; Harlan Laboratories, Boxmeer, The Netherlands) by microvascular photothrombosis as described previously [8]. This model mimics small arteries occlusion, commonly produced in human stroke, inducing a focal brain damage with histological and MRI correlations to human patterns [35]. Brain injury involves damage to vascular endothelium, activation of platelets, followed by microvascular thrombotic occlusion of a particular region [43], selected for irradiation by using a stereotaxic frame. In these experiments, we used coordinates +0.2 AP, +2 ML relative to Bregma reference point to damage the primary motor (hindlimb and forelimb) and somatosensory cortex according to the Paxinos mouse brain atlas. Thus, mice anesthetised with isoflurane (5% for induction, 2% for maintenance in oxygen, Abbot Laboratories, Madrid, Spain) were placed in a stereotaxic frame (Narishige Group, Tokyo, Japan), the body temperature being maintained at 36–37°C with a self-regulating heating blanket (Cibertec, Madrid, Spain). A midline scalp incision allowed to expose the skull and identify Bregma and Lambda points. Then, we used a micromanipulator to centre a cold-light (Schott KL 2500 LCD, Schott Glass, Mainz, Germany) with a fibre optic bundle of 1.5 mm in diameter at the indicated coordinates (right side). Next, the photosensitive dye Rose Bengal (20 mg/kg; Cat#R3877, Sigma-Aldrich) was administered by retro-orbital injection of the venous sinus, for intravenous (i.v.) vascular access, followed after 5 min by brain illumination through the intact skull for 10 min (600 lms, 3000K), leading to local dye activation and damage in those areas underneath the selected stereotaxic position. After completion of the surgical procedure, the incision was sutured and mice were allowed to recover.

As indicated, a single dose (10 nmol/g) of peptides MTMyc (Ac-YGRKKRRQRRRAEEQKLISEEDLLR-NH_2_), MTFL_457_ (Ac-YGRKKRRQRRRHSKFGMKGPASVIS-NH_2_) or MTFL_457_AAA (Ac-YGRKKRRQRRRHSAAAMKGPASVIS-NH_2_) (>95% purity; GenScript) were retro-orbitally injected 10 min after damage initiation, immediately after completing irradiation. These peptides are N-ter acetylated and C-ter amidated, for improvement of plasma stability. They were prepared as 2.5 mM solutions in 0.9% NaCl and, just before injection, acidity due to the trifluoracetic acid counterions present in the peptide preparations was neutralized by adding fresh HCO_3_NH_4_ to a final concentration of 44 mM. Mice were not subjected to other procedures before ischemia and were naïve to drug or peptide treatment. Animals were randomly allocated to the experimental groups and the researchers doing the experiments were blind respect to treatment. For immunohistochemistry, mice deeply anesthetized 5 h after damage were intracardially perfused with cold PBS and 4% paraformaldehyde in PBS. Brain processing proceeded as explained below. For assessment of infarct volume, animals were sacrificed by CO_2_ inhalation followed by cervical dislocation 24 h after damage induction. Their brains were sectioned into serial 1-mm-thick coronal slices using a mouse brain matrix (Stoelting, Wood Dale, IL, USA). Slices were completely stained with 2% TTC (Cat#T8877, Sigma-Aldrich) at room temperature, and fixed in 4% paraformaldehyde before scanning of both rostral and caudal sides.

### Primary culture of rat cortical neurons and treatment

Primary neuronal cultures were obtained from the brain cortex of 18-day-old Wistar rat embryos (E18), both genders being indistinctly used, as previously described (9]. Briefly, the dissected cortices were mechanically dissociated in Minimum Essential Medium supplemented with 22.2 mM glucose, 0.1 mM glutamax (Cat#35050-038, Gibco), 5% foetal bovine serum (Cat#FBS-HI-12A, Capricorn), 5% donor horse serum (Cat#11510516; Life Technologies), and 100 U/ml penicillin plus 100 µg/ml streptomycin (Cat#15140148; Life Technologies). Then, the cell suspension was seeded at a density of 1x10^6^ cells/ml in the same medium in plates previously treated with poly-L-lysine (100 µg/ml; Cat#P1524, Sigma-Aldrich) and laminin (4 µg/ml; Cat#L2020, Sigma-Aldrich) overnight at 37°C. These are mixed cultures, where early growth of the glial subpopulation helps to maintain non-toxic glutamate levels along neuronal maturation. After 7 days *in vitro* (DIVs), further glial proliferation was inhibited using cytosine ß-D-arabino furanoside (AraC, 10 µM; Cat#C1768, Sigma-Aldrich) and growth continued until 13 DIVs. Then, mature neurons were either subjected to excitotoxicity as below indicated or treated with BDNF (100 ng/ml; Cat#450-02, PeproTech) for the times specified. When indicated, cultures were preincubated with MTMyc, MTFL_457_ or MTFL_457_AAA (25 µM, 30 min) before excitotoxicity induction, peptides remaining in the culture media for the duration of the experiment. Mature neurons were also treated for 30 min with dynasore hydrate (80 μM; Cat#D7693, Sigma-Aldrich) before NMDA addition as indicated.

### Induction of neuronal excitotoxicity and evaluation of neuronal viability

Cultures were incubated for the indicated times with NMDA (100 µM; Cat#0114, Tocris) and its co-agonist glycine (10 µM; Cat#161-0718; Bio-Rad), herein denoted simply as NMDA, to induce a strong excitotoxic response in the mature neurons present in the mixed culture but no effect on astrocyte viability as previously described [64]. In order to analyse shedding of TrkB fragments due to MP action, culture medium was collected before cell lysis and analysed by immunoblot. Neuronal viability was measured by the MTT reduction assay. MTT (0.5 mg/ml; Cat#M5655, Sigma-Aldrich) was added to culture medium after 2 h of NMDA treatment and incubation proceeded for 2 additional hours at 37°C. Then, the produced formazan salts were solubilized in DMSO and spectrophotometrically quantified at 570 nm. In the primary cultures, the contribution of quiescent glial cells to the total values of cell viability was established by exposing sister cultures to 400 μM NMDA, 10 μM glycine for 24 h before the MTT assay. These conditions cause nearly complete death of mature neurons but no glial damage [64]. Once subtracted the absorbance value contributed by glial cells, we calculated the viability of the neuronal subpopulation in each sample. Every individual experiment included sample triplicates for each treatment, and multiple completely independent experiments were carried out as detailed in the figure legend. The viability of neurons preincubated with a peptide and subjected to excitotoxicity was calculated relative to that of neurons preincubated with the same peptide but no NMDA.

### Western blot analysis

Cultured cells were lysed in RIPA buffer (50 mM Tris-HCl pH 8, 150 mM NaCl, 1% sodium deoxycholate, 1% NP-40, 1 mM DTT and 0.1% SDS) with protease and phosphatase inhibitors (Complete protease and PhosSTOP phosphatases inhibitor cocktail tablets, Cat#11 697 498 001 and Cat#04 906 837 001, Roche). The concentration of protein was established with BCA Protein Assay Kit (Cat#A 2001, Thermo Fisher) and lysates were denatured in Laemmli SDS-sample buffer by heating at 95°C for 5 min. Equal amounts of total cell-lysates were resolved in Tris-Glycine SDS-PAGE and transferred on to a Protran nitrocellulose membrane (Cat#GE10600002, GE Healthcare). The efficacy of protein transfer to membranes was confirmed by staining with 1% (w/v) Ponceau S. After blocking with a 5% non-fat dry milk solution in Tris-buffered saline (TBS) with 0.05% Tween-20, membranes were incubated overnight at 4°C with primary antibodies, washed, and then incubated with appropriate anti-rabbit (Cat#A120-108P, RRID:AB_10892625; Bethyl) or anti-mouse (Cat#A90-137P; RRID:AB_1211460, Bethyl) peroxidase-conjugated secondary antibodies for 1 h at room temperature. Finally, immunoreactivity was visualized using Clarity Western ECL Blotting Substrate (Cat# 1705060, BioRad) and band intensity was quantified by densitometric analysis (Adobe Photoshop). Protein levels were normalized using those of neuron-specific enolase (NSE) present in the same sample and expressed relative to values obtained in their respective controls, arbitrarily given a 100% value. This neuronal loading control was chosen since NSE levels are not affected by NMDA treatment. In contrast, the activation of calpain induced by excitotoxicity was confirmed by analysing the formation of characteristic breakdown products (BDPs; 150 and 145 kDa) from brain spectrin, a standard substrate of this protease. Multiple and completely independent experiments were carried out and quantitated as detailed in the figure legends. Primary antibodies against the following proteins were used: Hrs (Cat#sc-271455, RRID:AB_10648901, Santa Cruz Biotechnology), NSE (Cat#AB951, RRID:AB_92390, Millipore), spectrin alpha chain (Cat#MAB1622, RRID:AB_94295, Millipore), TrkB-FL C-ter (sc-11, RRID:AB_632554, Santa Cruz Biotechnology), TrkB extracellular sequences or panTrkB (sc-136990, RRID:AB_2155262, Santa Cruz Biotechnology), PSD-95 C-ter (Cat#610496, RRID:AB_2315218, Transduction Laboratories), pY816-TrkB-FL (Cat#P01388, Boster).

### Protein immunoprecipitation

Total cell lysates were prepared from cortical cultures in 1% NP-40, 80 mM NaCl, 20 mM EDTA and 20 mM Tris-HCl (pH 8), containing protease and phosphatase inhibitors as before. Approximately 0.3-1 mg of cleared lysate was immunoprecipitated for 1 h at 4°C with 3-4 µg of antibody as indicated before the addition of 60 µl of 50% Protein G Agarose (Cat#15920010, Thermo Fisher) and incubation for 1 h at room temperature with agitation. After that, equivalent volumes of the immunoprecipitated complexes were analysed by immunoblot analysis as indicated and compared to equal amounts of the starting total lysates.

### Biotinylation of cell-surface proteins

After NMDA treatment for 1 h, cultures were immediately washed with ice-cold PBS containing 1 mM CaCl_2_ and 0.5 mM MgCl_2_ before biotinylation of surface proteins for 30 min at 4°C with 0.5 mg/ml of EZ-Link Sulfo-NHS-SS-biotin (Cat#21331, Thermo Fisher) prepared in that same buffer. The excess of free biotin was removed by washing cultures with cold PBS containing 1 mM CaCl_2_, 0.5 mM MgCl_2_, and 0.1% BSA, followed by two additional washes without BSA. Next, cells were lysed in RIPA buffer containing protease and phosphatase inhibitors but without DTT, a small aliquot of the total extracts being saved for further analysis. The remaining total extracts were incubated with streptavidin resin (Cat#L00353, GenScript) for 3 h at 4°C to precipitate the biotinylated proteins. After washing the streptavidin–biotin complexes twice with lysis buffer containing 500 mM NaCl, protease and phosphatase inhibitors, plus two additional washing steps omitting this salt, pellets were solubilized and denature in SDS–PAGE sample buffer (10 min at 50°C). Equivalent volumes of the isolated proteins and equal amounts of the total protein extracts were analysed in parallel by Western blot. Totally independent experiments were carried out and quantified as detailed in figure legends.

### Immunocytochemistry

Primary cultures grown as before on coverslips coated with poly-L-lysine/laminin were treated as indicated and fixed with 4% paraformaldehyde in PBS for 30 min. After washing with PBS, cells were blocked and permeabilized for 30 min at room temperature with 1% BSA, 0.1% Triton X-100 in PBS. Afterwards coverslips were incubated overnight at 4°C with primary antibodies diluted in the same solution recognizing the following epitopes: Hrs (Cat#sc-271455, RRID:AB_10648901, Santa Cruz Biotechnology), TrkB-FL C-ter (sc-11, RRID:AB_632554, Santa Cruz Biotechnology) and GM130 (Cat# A-610822, RRID:AB_398141, BD Biosciences). Detection was achieved with goat secondary antibodies conjugated to Alexa Fluor 488 (anti-rabbit, Cat #A-11034, RRID:AB_2576217; anti-mouse, Cat#A-11029, RRID:AB_ 2534088, Thermo Fisher Scientific) or Alexa Fluor 546 (anti-rabbit, Cat#A-11035, RRID:AB_ 2534093, Thermo Fisher Scientific), diluted as above. After that, coverslips were incubated 10 min in a 0.5 µg/ml of DAPI (Cat#D1306, Molecular Probes) before mounting with Prolong Diamond antifade reagent (Cat#P36970, Molecular Probes). All confocal images are single sections acquired using an inverted Zeiss LSM 710 laser confocal microscope (Jena, Germany) with a 63x Plan-Apochromatic oil immersion objective and were normalized for each colour separately. Images were processed for presentation with ImageJ (NIH Image).

For colocalization studies, we analysed a minimum of 80 neurons per experiment and quantified four independent experiments. We calculated the Pearson correlation coefficient (PCC) as well as the percentage of cells having PCC values ≥ 0.50 (or the indicated threshold). To analyse GA fragmentation, we used the Fiji image processing package (https://www.nature.com/articles/nmeth.2019; RRID:SCR_002285) and a macro established by Dr. Sánchez-Ruiloba (Instituto de Investigaciones Marinas, CSIC). All images were analysed in a similar way. Among optical sections acquired in the confocal microscope, we selected a central one where labelled particles were best detected and, after establishing a threshold level to consider a signal as positive, we identified the particles and obtained for each of them the values of area, mean intensity, circularity and integrated density. We analysed 3-5 images corresponding to the different conditions in each experiment, and calculated the mean values of 5 independent experiments. We also chose the image that best fitted the final mean values for representing the specific values obtained for each particle in a particular condition and experiment.

### Measurement of infarct volume

Rostral and caudal images of coronal slices stained with TTC as described were analysed using ImageJ software (https://imagej.net/; RRID:SCR_003070) by an observer blinded to experimental groups. After image calibration, delineated areas of ipsilateral and contralateral hemispheres, and the infarcted region (unstained area) were measured. Considering slices thickness, the corresponding volumes were calculated and corrected for oedema’s effect, estimated by comparing total volumes of hemispheres. The corrected infarct volumes were expressed as percentage relative to the contralateral hemisphere, to correct for normal size differences between different animals. For each animal, the mean of results obtained for rostral and caudal sides was calculated.

### Immunohistochemistry and Fluoro-Jade C staining

Brains obtained from animals sacrificed 5 h after injury were post-fixed in 4% paraformaldehyde in PBS at 4°C for 24 h and cryoprotected in 30% sucrose for 48 h at 4°C. Coronal frozen sections (30 μm thick) were prepared using a cryostat (Leica, Heidelberg, Germany) and sections were incubated with blocking solution (10% goat serum, 0.5% Triton X-100 in PB) in flotation for 1 h at room temperature and then overnight at 4°C with the mouse antibody that recognizes protein GM130 (Cat# A-610822, RRID:AB_398141, BD Biosciences), followed by incubation for 2 h at room temperature with Alexa Fluor 546-conjugated goat anti-mouse secondary antibodies (Cat#A-11030, RRID:AB_ 2534089, Thermo Fisher Scientific) and DAPI (5 µg/ml; Cat#D1306, Molecular Probes) to stain DNA. After washing in distilled water, sections were mounted on slides, dried overnight at RT, cleared in xylene and then cover slipped with DPX (Cat#06522, Sigma-Aldrich). As a control experiment, incubation with the anti-GM130 antibody was omitted when indicated. Additionally, in the indicated experiments, blocking was followed by incubation for 18 h at 4°C with a goat anti-mouse IgG H&L antibody (Fab fragment; Cat#ab6668, RRID:AB_955960, Abcam) diluted in 5% goat serum, 0.5% Triton X-100 in PB, before washing and incubation with the secondary antibody. This additional step decreased the background due to reaction of anti-mouse secondary antibodies with mouse antibodies leaked to the ischemic area as a consequence of early BBB breakage induced by damage [46).

For combined double immunohistochemistry and Fluoro-Jade C staining, after blockage, brain sections were incubated overnight at 4°C with mouse anti-GM130 and rabbit anti-SNTF antibodies (Cat#ABN2264, Millipore) followed by incubation for 2 h at room temperature with goat Alexa Fluor 546-conjugated anti-mouse (Cat#A-11030; RRID:AB_2534089, Thermo Fisher Scientific) and Alexa Fluor 647-conjugated anti-rabbit secondary antibodies (Cat#A-21245, RRID:AB_2535813, Thermo Fisher Scientific). Next, sections were rinsed in distilled water and incubated with a solution of 0.0004% Fluoro-Jade C (Cat#AG325, Millipore) and 0.0002% DAPI in 0.01% acetic acid for 10 min at room temperature for specific labelling of, respectively, degenerating neurons of the infarcted area and cell nuclei. Unless otherwise indicated, confocal images were acquired with a 63x Plan-Apochromatic oil immersion objective and processed for presentation as above described.

### Quantification and statistical analysis

All data were expressed as mean ± standard error of the mean (SEM) of 3 to 9 completely independent experiments. The details of the number of experiments done, the precise sample size and the specific statistical test applied for each case can be found in the figure legends. Cell samples in each individual experiment were sister primary cultures grown in multiwell plates, obtained from the same cell suspension, treatments being assigned in a random way. Statistical analysis was performed with the Statistical Package for Social Science (SPSS, v.18, RRID:SCR_002865, IBM) and GraphPad Prism 8.4.3.686 (https://www.graphpad.com; RRID:SCR_002798). Considering the number of groups, the data distribution and the homogeneity of variances, we used unpaired Student’s t-test or one-way ANOVA followed by Bonferroni honestly significant difference test (HSD) or a multiple comparison test (LSD *post-hoc* test). Data were represented as per cent of controls or maximum values as indicated. A *P* value smaller than 0.05 was considered statistically significant (*\*P* < 0.05, \**\* P* < 0.01, \*\**\* P* < 0.001).

## Acknowledgements

We are grateful to Dr. Lucía Sánchez-Ruiloba (Microscopy and Image Analysis Unit, Instituto de Investigaciones Marinas, CSIC, Spain) for technical advice with the quantitation of GA fragmentation and Dr. Esther San Antonio (Immunology Department, Instituto de Investigación Sanitaria-Princesa, Spain) for her help with initial experiments in this project. We also thank members of our group for helpful discussions. The authors declare no competing financial interests.

## Authors’ Contributions

GME-O designed and performed experiments, analysed data, and reviewed/edited the manuscript draft; MD-G secured funding, designed experiments and supervised project, contributed to formal analysis, wrote the original draft, and reviewed and edited the manuscript. All authors read and approved the final manuscript.

## Ethics approval

This study has no human data. Animal studies were approved by ethics committees from Consejo Superior de Investigaciones Científicas (CSIC) and Comunidad de Madrid (Ref PROEX 276.6/20).

## Data availability

All data supporting the conclusions of this article are included within the article and in the additional file provided.

---

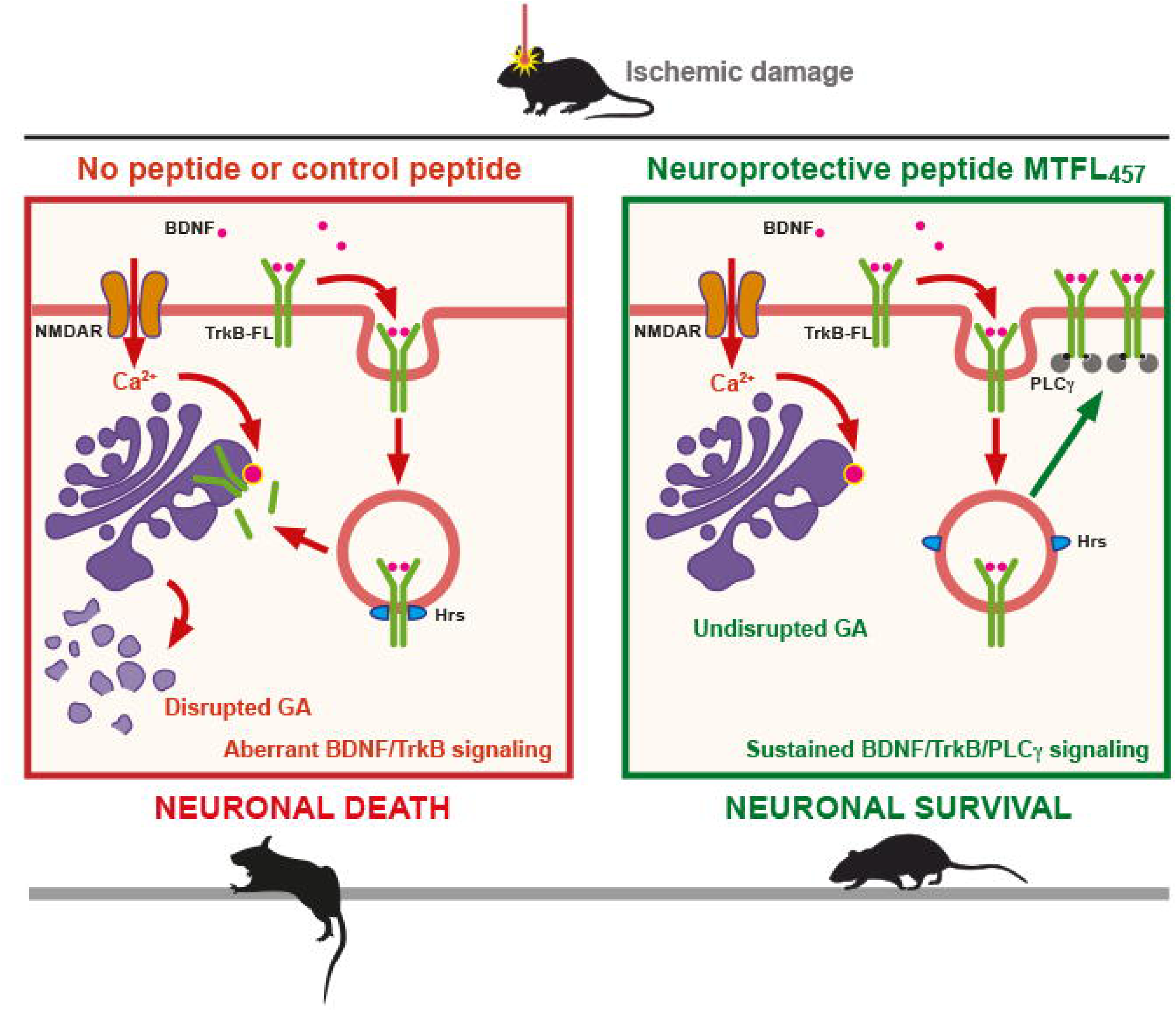

## References

1. Choi DW. Excitotoxicity: Still Hammering the Ischemic Brain in 2020. Front Neurosci. 2020;14:579953.

2. Choi DW. Glutamate neurotoxicity and diseases of the nervous system. Neuron. 1988;1:623–34.

3. Wu Q & Tymianski M. Targeting NMDA receptors in stroke: new hope in neuroprotection. Mol Brain. 2018;11:15.

4. Zhou XF. ESCAPE-NA1 Trial Brings Hope of Neuroprotective Drugs for Acute Ischemic Stroke: Highlights of the Phase 3 Clinical Trial on Nerinetide. Neurosci Bull. 2021;37:579–81.

5. Rex N, Ospel JM, McDonough RV, Kashani N, Rinkel L, Buck BH, et al. Nerinetide reduces early infarct growth among stroke patients undergoing EVT without intravenous alteplase. Stroke: Vasc Interv Neurol. 2024; 4:e001034.

6. Hill MD, Martin RH, Mikulis D, Wong JH, Silver FL, Terbrugge KG, et al. Safety and efficacy of NA-1 in patients with iatrogenic stroke after endovascular aneurysm repair (ENACT): a phase 2, randomised, double-blind, placebo-controlled trial. Lancet Neurol. 2012;11:942–50.

7. Kurrikoff K & Langel U. Recent CPP-based applications in medicine. Expert Opin Drug Deliv. 2019;16:1183–91.

8. Tejeda GS, Esteban-Ortega GM, San Antonio E, Vidaurre ÓGG & Díaz-Guerra M. Prevention of excitotoxicity-induced processing of BDNF receptor TrkB-FL leads to stroke neuroprotection. EMBO Mol Med. 2019, 11:e9950.

9. Ayuso-Dolado S, Esteban-Ortega GM, Vidaurre OG & Diaz-Guerra M. A novel cell-penetrating peptide targeting calpain-cleavage of PSD-95 induced by excitotoxicity improves neurological outcome after stroke. Theranostics. 2021;11:6746–65.

10. Reichardt LF. Neurotrophin-regulated signalling pathways. Philos Trans R Soc Lond B Biol Sci. 2006;361:1545–64.

11. Huang EJ & Reichardt LF. Trk receptors: roles in neuronal signal transduction. Annu Rev Biochem. 2003;72:609–42.

12. Bonni A, Brunet A, West AE, Datta SR, Takasu MA & Greenberg ME. Cell survival promoted by the Ras-MAPK signaling pathway by transcription-dependent and -independent mechanisms. Science. 1999;286:1358–62.

13. Liu L, Cavanaugh JE, Wang Y, Sakagami H, Mao Z & Xia Z. ERK5 activation of MEF2-mediated gene expression plays a critical role in BDNF-promoted survival of developing but not mature cortical neurons. Proc Natl Acad Sci U S A. 2003;100:8532–7.

14. Wang Y, Liu L & Xia Z. Brain-derived neurotrophic factor stimulates the transcriptional and neuroprotective activity of myocyte-enhancer factor 2C through an ERK1/2-RSK2 signaling cascade. J Neurochem. 2007;102:957–66.

15. Kingsbury TJ, Murray PD, Bambrick LL & Krueger BK. Ca(2+)-dependent regulation of TrkB expression in neurons. J Biol Chem. 2003;278:40744–8.

16. Deogracias R, Espliguero G, Iglesias T & Rodriguez-Pena A. Expression of the neurotrophin receptor trkB is regulated by the cAMP/CREB pathway in neurons. Mol Cell Neurosci. 2004;26:470–80.

17. Tao X, Finkbeiner S, Arnold DB, Shaywitz AJ & Greenberg ME. Ca2+ influx regulates BDNF transcription by a CREB family transcription factor-dependent mechanism. Neuron. 1998;20:709–26.

18. Shieh PB, Hu SC, Bobb K, Timmusk T & Ghosh A. Identification of a signaling pathway involved in calcium regulation of BDNF expression. Neuron. 1998;20:727–40.

19. Lyons MR, Schwarz CM & West AE. Members of the myocyte enhancer factor 2 transcription factor family differentially regulate Bdnf transcription in response to neuronal depolarization. J Neurosci. 2012;32:12780–5.

20. Haapasalo A, Koponen E, Hoppe E, Wong G & Castren E. Truncated trkB.T1 is dominant negative inhibitor of trkB.TK+-mediated cell survival. Biochem Biophys Res Commun. 2001;280:1352–8.

21. Zheng J, Shen WH, Lu TJ, Zhou Y, Chen Q, Wang Z, et al. Clathrin-dependent endocytosis is required for TrkB-dependent Akt-mediated neuronal protection and dendritic growth. J Biol Chem. 2008;283:13280–8.

22. Huang SH, Zhao L, Sun ZP, Li XZ, Geng Z, Zhang KD, et al. Essential role of Hrs in endocytic recycling of full-length TrkB receptor but not its isoform TrkB.T1. J Biol Chem. 2009;284:15126–36.

23. Berretta A, Tzeng Y-C & Clarkson AN. Post-stroke recovery: the role of activity-dependent release of brain-derived neurotrophic factor. Expert Rev Neurother. 2014;14:1335–44.

24. Tejeda GS & Diaz-Guerra M. Integral characterization of defective BDNF/TrkB signalling in neurological and psychiatric disorders leads the way to new therapies. Int J Mol Sci. 2017;18:e268.

25. Vidaurre OG, Gascón S, Deogracias R, Sobrado M, Cuadrado E, Montaner J, et al. Imbalance of neurotrophin receptor isoforms TrkB-FL/TrkB-T1 induces neuronal death in excitotoxicity. Cell Death & Dis. 2012;3:e256.

26. Tejeda GS, Ayuso-Dolado S, Arbeteta R, Esteban-Ortega GM, Vidaurre OG & Diaz-Guerra M. Brain ischaemia induces shedding of a BDNF-scavenger ectodomain from TrkB receptors by excitotoxicity activation of metalloproteinases and gamma-secretases. J Pathol. 2016;238:627–40.

27. Fonseca-Gomes J, Jerónimo-Santos A, Lesnikova A, Casarotto P, Castrén E, Sebastião AM, et al. TrkB-ICD fragment, originating from BDNF receptor cleavage, is translocated to cell nucleus and phosphorylates nuclear and axonal proteins. Front Mol Neurosci. 2019;12:4.

28. Gonatas NK, Stieber A & Gonatas JO. Fragmentation of the Golgi apparatus in neurodegenerative diseases and cell death. J Neurol Sci. 2006;246:21–30.

29. Park JS, Bateman MC & Goldberg MP. Rapid alterations in dendrite morphology during sublethal hypoxia or glutamate receptor activation. Neurobiol Dis. 1996;3:215–27.

30. Vosler PS, Brennan CS & Chen J. Calpain-mediated signaling mechanisms in neuronal injury and neurodegeneration. Mol Neurobiol. 2008;38:78–100.

31. Vaslin A, Puyal J, Borsello T & Clarke PG. Excitotoxicity-related endocytosis in cortical neurons. J Neurochem. 2007;102:789–800.

32. Vaslin A, Puyal J & Clarke PG. Excitotoxicity-induced endocytosis confers drug targeting in cerebral ischemia. Ann Neurol. 2009;65:337–47.

33. Geetha T, Jiang J & Wooten MW. Lysine 63 polyubiquitination of the nerve growth factor receptor TrkA directs internalization and signaling. Mol Cell. 2005;20:301–12.

34. Guo Y-YY, Lu Y, Zheng Y, Chen X-RR, Dong J-LL, Yuan R-RR, et al. Ubiquitin C-terminal hydrolase L1 (UCH-L1) promotes hippocampus-dependent memory via its deubiquitinating effect on TrkB. J Neurosci. 2017;37:5978–95.

35. Pevsner PH, Eichenbaum JW, Miller DC, Pivawer G, Eichenbaum KD, Stern A, et al. A photothrombotic model of small early ischemic infarcts in the rat brain with histologic and MRI correlation. J Pharmacol Toxicol Methods. 2001;45:227–33.

36. Schreij AM, Fon EA & McPherson PS. Endocytic membrane trafficking and neurodegenerative disease. Cell Mol Life Sci. 2016;73:1529–45.

37. Komada M, Masaki R, Yamamoto A & Kitamura N. Hrs, a tyrosine kinase substrate with a conserved double zinc finger domain, is localized to the cytoplasmic surface of early endosomes. J Biol Chem. 1997;272:20538–44.

38. Lazo OM, Gonzalez A, Ascano M, Kuruvilla R, Couve A & Bronfman FC. BDNF regulates Rab11- mediated recycling endosome dynamics to induce dendritic branching. J Neurosci. 2013;33:6112–22.

39. Gamir-Morralla A, López-Menéndez C, Ayuso-Dolado S, Tejeda GS, Montaner J, Rosell A, et al. Development of a neuroprotective peptide that preserves survival pathways by preventing Kidins220/ARMS calpain processing induced by excitotoxicity. Cell Death & Dis. 2015;6:e1939.

40. López-Menéndez C, Simón-García A, Gamir-Morralla A, Pose-Utrilla J, Luján R, Mochizuki N, et al. Excitotoxic targeting of Kidins220 to the Golgi apparatus precedes calpain cleavage of Rap1- activation complexes. Cell Death & Dis. 2019;10:535.

41. Rajagopal R, Chen ZY, Lee FS & Chao MV. Transactivation of Trk neurotrophin receptors by G-protein-coupled receptor ligands occurs on intracellular membranes. J Neurosci. 2004;24:6650–8.

42. Nakagomi S, Barsoum MJ, Bossy-Wetzel E, Sutterlin C, Malhotra V & Lipton SA. A Golgi fragmentation pathway in neurodegeneration. Neurobiol Dis. 2008;29:221–31.

43. Schroeter M, Jander S & Stoll G. Non-invasive induction of focal cerebral ischemia in mice by photothrombosis of cortical microvessels: characterization of inflammatory responses. J Neurosci Methods. 2002;117:43–9.

44. Carmichael ST. Rodent models of focal stroke: size, mechanism, and purpose. NeuroRx. 2005;2:396–409.

45. Yuan D, Hu K, Loke CM, Teramoto H, Liu C & Hu B. Interruption of endolysosomal trafficking leads to stroke brain injury. Exp Neurol. 2021;345:113827.

46. Abdullahi W, Tripathi D & Ronaldson PT. Blood-brain barrier dysfunction in ischemic stroke: targeting tight junctions and transporters for vascular protection. Am J Physiol Cell Physiol. 2018;315:C343–C56.

47. Sutterlin C, Hsu P, Mallabiabarrena A & Malhotra V. Fragmentation and dispersal of the pericentriolar Golgi complex is required for entry into mitosis in mammalian cells. Cell. 2002;109:359–69.

48. Thayer DA, Jan YN & Jan LY. Increased neuronal activity fragments the Golgi complex. Proc Natl Acad Sci U S A. 2013;110:1482–7.

49. Joshi G, Bekier ME & Wang Y. Golgi fragmentation in Alzheimer’s disease. Front Neurosci. 2015;9:340.

50. Mizuno Y, Hattori N, Kitada T, Matsumine H, Mori H, Shimura H, et al. Familial Parkinson’s disease. Alpha-synuclein and parkin. Adv Neurol. 2001;86:13–21.

51. Hilditch-Maguire P, Trettel F, Passani LA, Auerbach A, Persichetti F & MacDonald ME. Huntingtin: an iron-regulated protein essential for normal nuclear and perinuclear organelles. Hum Mol Genet. 2000;9:2789–97.

52. Fujita Y & Okamoto K. Golgi apparatus of the motor neurons in patients with amyotrophic lateral sclerosis and in mice models of amyotrophic lateral sclerosis. Neuropathology. 2005;25:388–94.

53. Schecterson LC, Hudson MP, Ko M, Philippidou P, Akmentin W, Wiley J, et al. Trk activation in the secretory pathway promotes Golgi fragmentation. Mol Cell Neurosci. 2010;43:403–13.

54. Wu Y, Chen C, Yang Q, Jiao M & Qiu S. Endocytosis of GluN2B-containing NMDA receptors mediates NMDA-induced excitotoxicity. Mol Pain. 2017;13:1744806917701921.

55. Gascon S, Sobrado M, Roda JM, Rodriguez-Pena A & Diaz-Guerra M. Excitotoxicity and focal cerebral ischemia induce truncation of the NR2A and NR2B subunits of the NMDA receptor and cleavage of the scaffolding protein PSD-95. Mol Psychiatry. 2008;13:99–114.

56. Lopez-Menendez C, Gascon S, Sobrado M, Vidaurre OG, Higuero AM, Rodriguez-Pena A, et al. Kidins220/ARMS downregulation by excitotoxic activation of NMDARs reveals its involvement in neuronal survival and death pathways. J Cell Sci. 2009;122:3554–65.

57. Tompa P, Emori Y, Sorimachi H, Suzuki K & Friedrich P. Domain III of calpain is a ca2+-regulated phospholipid-binding domain. Biochem Biophys Res Commun. 2001;280:1333–9.

58. Hood JL, Brooks WH & Roszman TL. Differential compartmentalization of the calpain/calpastatin network with the endoplasmic reticulum and Golgi apparatus. J Biol Chem. 2004;279:43126–35.

59. Hood JL, Brooks WH & Roszman TL. Subcellular mobility of the calpain/calpastatin network: an organelle transient. Bioessays. 2006;28:850–9.

60. Bevers MB & Neumar RW. Mechanistic role of calpains in postischemic neurodegeneration. J Cereb Blood Flow Metab. 2008;28:655–73.

61. Diaz-Guerra M. Excitotoxicity-induced endocytosis as a potential target for stroke neuroprotection. Neural Regen Res. 2021;16:300–1.

62. Casarotto PC, Girych M, Fred SM, Kovaleva V, Moliner R, Enkavi G, et al. Antidepressant drugs act by directly binding to TRKB neurotrophin receptors. Cell. 2021;184:1299–313 e19.

63. Rantamaki T, Hendolin P, Kankaanpaa A, Mijatovic J, Piepponen P, Domenici E, et al. Pharmacologically diverse antidepressants rapidly activate brain-derived neurotrophic factor receptor TrkB and induce phospholipase-Cgamma signaling pathways in mouse brain. Neuropsychopharmacology. 2007;32:2152–62.

64. Choi DW, Maulucci-Gedde M & Kriegstein AR. Glutamate neurotoxicity in cortical cell culture. J Neurosci. 1987;7:357–68.

