## Supplementary figures for "Retrograde transport of neurotrophin receptor TrkB-FL induced by excitotoxicity regulates Golgi stability and is a target for stroke neuroprotection"

Fig. S1 Analysis of a possible effect of excitotoxicity on Hrs levels

Fig. S2 Effect of excitotoxicity on TrkB-FL/Hrs coimmunoprecipitation.

Fig. S3 Regulation by peptide MTFL457 of excitotoxicity-induced GA fragmentation.

Fig. S4 Leakage of mouse immunoglobulins to the brain cortex due to BBB breakage.

Fig. S5 Leakage of mouse immunoglobulins causes high backgrounds in immunohistochemistry of the ischemic tissue.

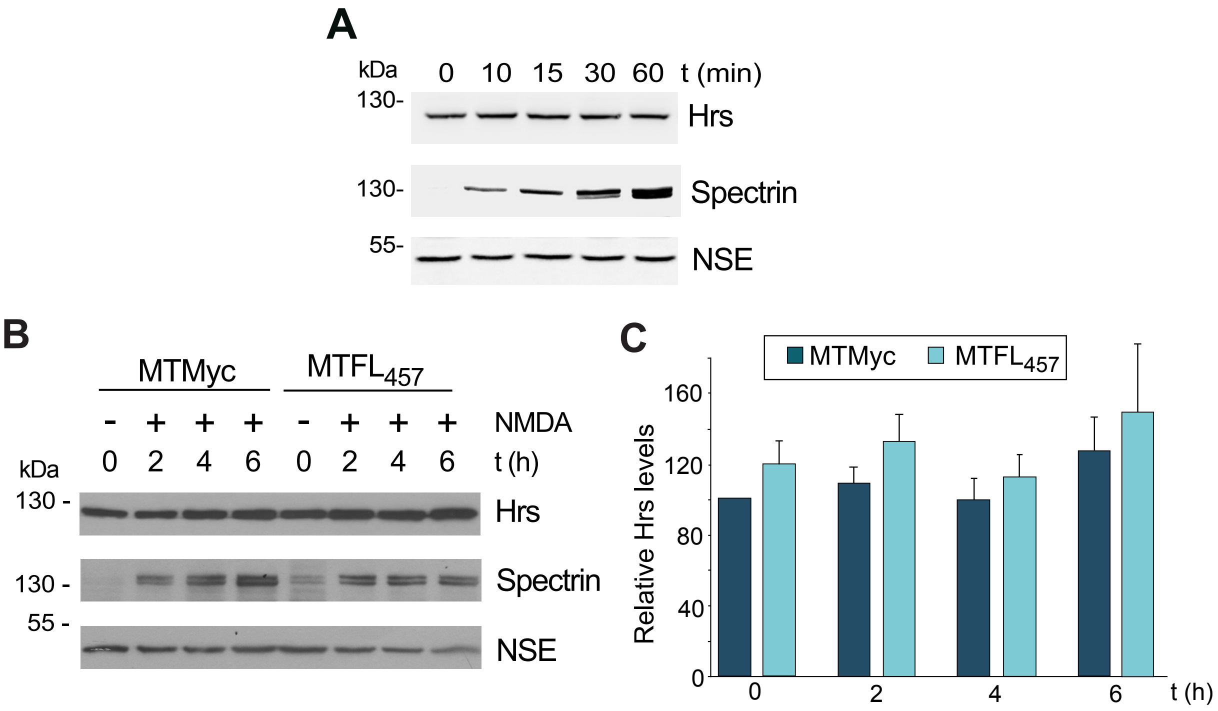


**Fig. S1 Analysis of a possible effect of excitotoxicity on Hrs levels. A** Cortical neurons were briefly treated with NMDA (0-60 min) and levels of endosomal protein Hrs established by immunoblot. Calpain activation was demonstrated by cleavage of spectrin. **B** Cell cultures were preincubated with MTMyc and $MTFL_{457}$ (25 μM, 30 min) and subsequently treated with NMDA for the indicated times (0-6 h). Levels of Hrs were analyze as above. **C** Mean ± SEM (n = 11) of normalized Hrs levels relative to those obtained in MTMyc-preincubated neurons in the absence of NMDA. Statistical analysis was performed using a generalized linear model followed by a *post-hoc* LSD test and no statistically significant differences were found.

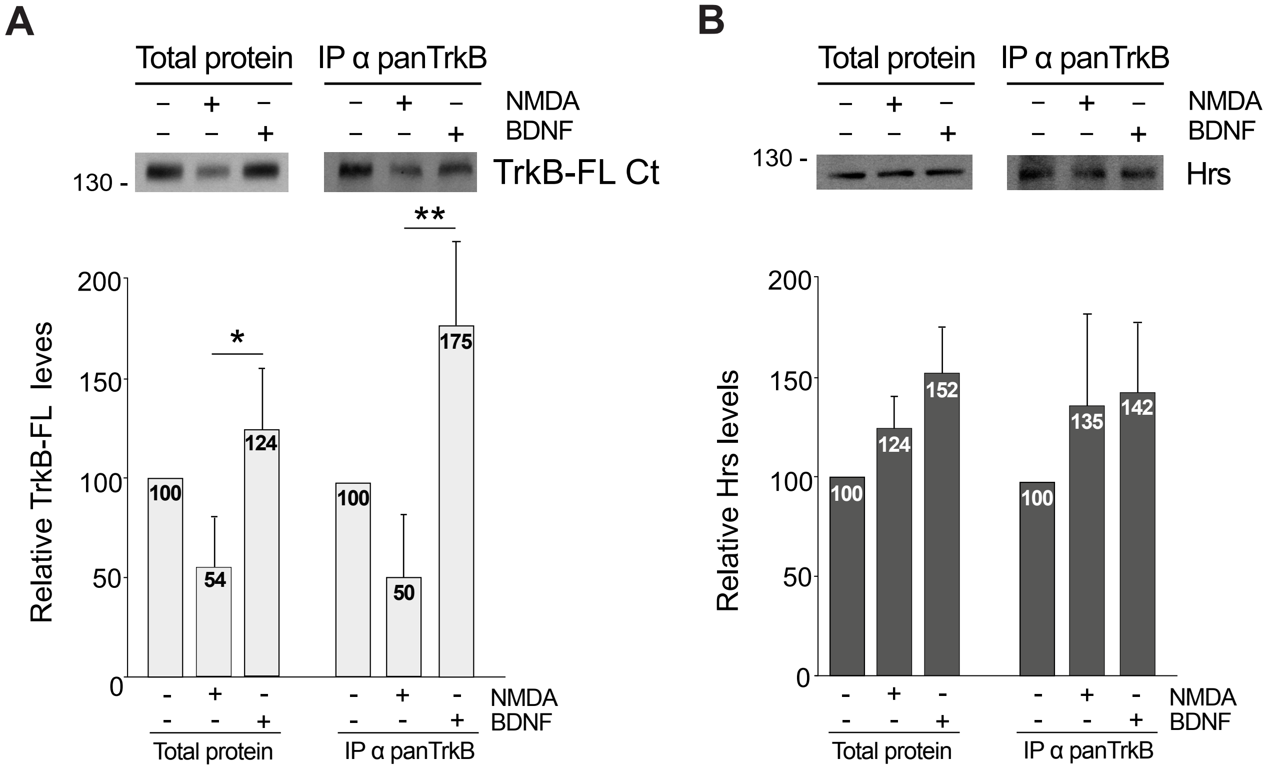


**Fig. S2 Effect of excitotoxicity on TrkB-FL/Hrs coimmunoprecipitation.** Neuronal cultures were treated with NMDA (100 µM) or BDNF (100 ng/ml) for 30 min and compared to untreated cultures. Proteins immunoprecipitated with antibody panTrkB (IP) were analyzed by immunoblot with the TrkB-FL Ct antibody **(A)** or that recognizing Hrs **(B)** in parallel to the corresponding total protein lysates. Mean values ± SEM (n = 5) of TrkB-FL and Hrs levels in NMDA or BDNF-treated cultures relative to untreated cells is represented. Statistical analysis was performed using a generalized linear model followed by a *post-hoc* LSD test (*$P < 0.05$, **$P < 0.01$).

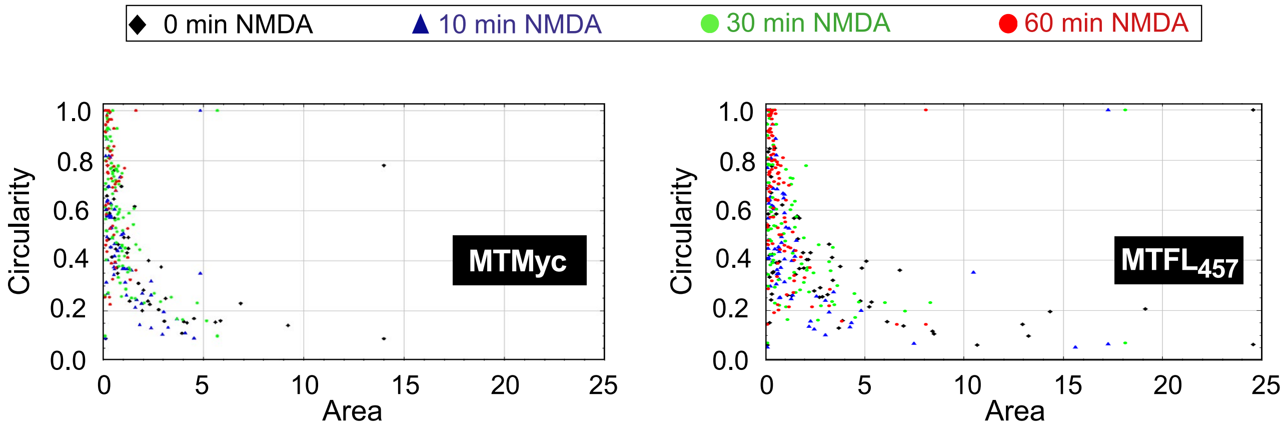


**Fig. S3 Regulation by peptide MTFL$_{457}$ of excitotoxicity-induced GA fragmentation.** Cortical neurons preincubated with MTMyc and MTFL$_{457}$ (25 µM, 30 min) and treated with NMDA for 0-60 min were analyzed by immunofluorescence with the GM130 antibody to visualize GA disruption as described. Representation of area versus circularity for each of the GA particles detected in representative images corresponding to cultures preincubated with MTMyc (left panel) or MTFL$_{457}$ (right panel), and treated with NMDA for the indicated times. A 0 value indicates absence of circularity, and 1 total circularity.

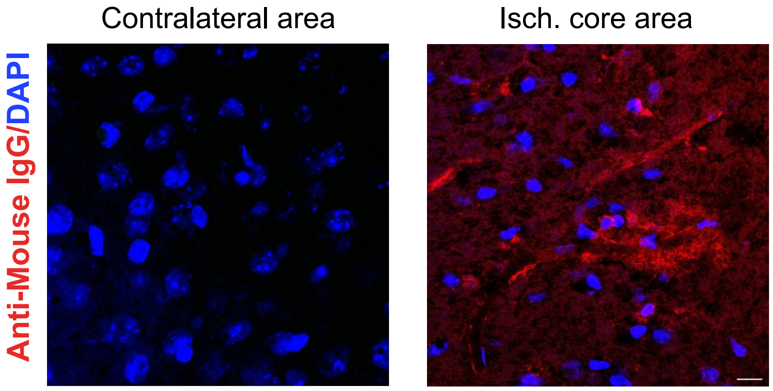


**Fig. S4 Leakage of mouse immunoglobulins to the brain cortex due to BBB breakage.** Brain coronal sections of animals sacrificed 5 h after insult were analyzed by immunohistochemistry with an anti-mouse secondary antibody, without the primary antibody. Heavy staining of blood vessels and high backgrounds, which challenge GM130 detection, were specifically observed in the ischemic tissue. Representative images correspond to single sections. Scale bar: 10 µm.

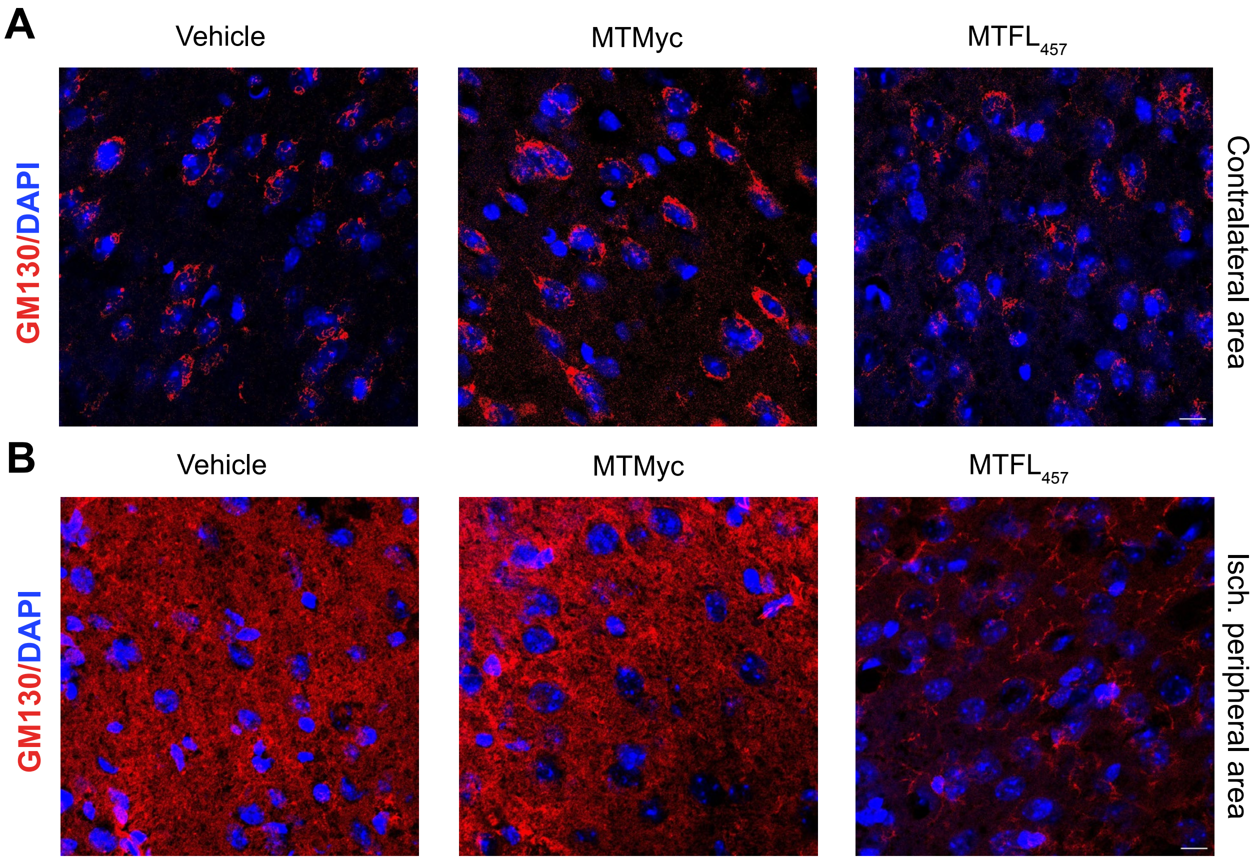


**Fig. S5 Leakage of mouse immunoglobulins causes high backgrounds in immunohistochemistry of the ischemic tissue.** Animals retro-orbitally injected with peptides MTMyc or $MTFL_{457}$ (10 nmol/g) or vehicle 10 min after damage initiation were sacrificed 5 h later. Comparison of GM130 staining in the contralateral **(A)** or the ischemic peripheral area **(B)** is shown. Representative images correspond to single sections. Scale bar: 10 µm.
